# Deep-diffeomorphic networks for conditional brain templates

**DOI:** 10.1101/2024.07.05.602288

**Authors:** Luke Whitbread, Stephan Laurenz, Lyle J Palmer, Mark Jenkinson, the Alzheimer’s Disease Neuroimaging Initiative

**Author notes:** Data used in preparation of this article were obtained from the Alzheimer’s Disease Neuroimaging Initiative (ADNI) database (adni.loni.usc.edu). As such, the investigators within the ADNI contributed to the design and implementation of ADNI and/or provided data but did not participate in analysis or writing of this report. A complete listing of ADNI investigators can be found at: http://adni.loni.usc.edu/wp-content/uploads/how_to_apply/ADNI_Acknowledgement_List.pdf. {; }. **Data Availability Statement:** The data that support the findings of this study are available from the Alzheimer’s Disease Neuroimaging Initiative (ADNI) database. Access to this data is subject to ADNI data sharing policies and restrictions. **Ethics Approval Statement:** As per ADNI protocols, all procedures performed in studies involving human participants were in accordance with the ethical standards of the institutional and/or national research committee and with the 1964 Helsinki declaration and its later amendments or comparable ethical standards. More details can be found at adni.loni.usc.edu. (This article does not contain any studies with human participants performed by any of the authors). **Patient Consent Statement:** Written informed consent was obtained from all participants included in the study by the ADNI investigators. **Permission to Reproduce Material:** Not applicable.

## Abstract

Deformable brain templates are an important tool in many neuroimaging analyses. Conditional templates (e.g., age-specific templates) have advantages over single population templates by enabling improved registration accuracy and capturing common processes in brain development and degeneration. Conventional methods require large, evenly-spread cohorts to develop conditional templates, limiting their ability to create templates that could reflect richer combinations of clinical and demographic variables. More recent deep-learning methods, which can infer relationships in very high dimensional spaces, open up the possibility of producing conditional templates that are jointly optimised for these richer sets of conditioning parameters. We have built on recent deep-learning template generation approaches using a diffeomorphic (topology-preserving) framework to create a purely geometric method of conditional template construction that learns diffeomorphisms between: (i) a global or group template and conditional templates, and (ii) conditional templates and individual brain scans. We evaluated our method, as well as other recent deep-learning approaches, on a dataset of cognitively normal (**CN**) participants from the Alzheimer’s Disease Neuroimaging Initiative (ADNI), using age as the conditioning parameter of interest. We assessed the effectiveness of these networks at capturing age-dependent anatomical differences. Our results demonstrate that while the assessed deep-learning methods have a number of strengths, they require further refinement to capture morphological changes in ageing brains with an acceptable degree of accuracy. The volumetric output of our method, and other recent deep-learning approaches, across four brain structures (grey matter, white matter, the lateral ventricles and the hippocampus), was measured and showed that although each of the methods captured some changes well, each method was unable to accurately track changes in all of the volumes. However, as our method is purely geometric it was able to produce T1-weighted conditional templates with high spatial fidelity and with consistent topology as age varies, making these conditional templates advantageous for spatial registrations. The use of diffeomorphisms in these deep-learning methods represents an important strength of these approaches, as they can produce conditional templates that can be explicitly linked, geometrically, across age as well as to fixed, unconditional templates or brain atlases. The use of deep-learning in conditional template generation provides a framework for creating templates for more complex sets of conditioning parameters, such as pathologies and demographic variables, in order to facilitate a broader application of conditional brain templates in neuroimaging studies. This can aid researchers and clinicians in their understanding of how brain structure changes over time, and under various interventions, with the ultimate goal of improving the calibration of treatments and interventions in personalised medicine. The code to implement our conditional brain template network is available at: github.com/lwhitbread/deep-diff.

## 1 Introduction

Magnetic Resonance Imaging (MRI) has evolved into an indispensable tool for the study of the human brain, providing insights into its structural and functional architecture without the need for invasive procedures [1]. For instance, as the brain ages, it undergoes various structural changes, such as reductions in grey-matter volume [2]. MRI’s ability to provide clear, detailed and differentiated images of soft tissues makes it well suited for obtaining insights into ageing processes in the brain and the progression of neurodegenerative diseases.

Deformable templates are reference images that can undergo geometric deformations to match participant scans contained in a dataset, and hence allow analyses of geometric variability [3]. Templates are utilised in a number of fields, including: (i) computer vision [4, 5, 6], (ii) graphics [7, 8], and medical image analysis [9, 10, 11, 12]. Brain templates and atlases are important tools for neuroimaging analyses, with applications including the characterisation of structural changes across the lifespan and the identification of pathological changes in the brain [13]. Conditional brain templates are tailored to reflect a particular set of conditioning parameters and have an important role in characterising changes in the brain that are associated with various factors such as age, environmental exposures and a range of other demographic, epidemiological, and clinical variables [14].

Conditional templates can aid researchers and clinicians in a variety of neuroimaging analyses, such as understanding how brain structure changes over time, under various interventions, or as disease progresses [15, 16]. By examining various structural measures of interest, including how volumes of specific brain regions (e.g., hippocampus, ventricles) vary as clinical and demographic variables change, researchers and clinicians can gain insight into the patterns of brain development and degeneration [17]. In research settings, it is also useful to have unbiased templates for studying older populations. These templates should be geometrically linked to a common reference template, to which standard atlases are also related. Such linkage is an essential facilitator of anatomical mapping across different templates, especially since it is not always feasible to create study-specific atlases.

Template construction methods have been studied for decades, undergoing many technical developments [18, 3]. These methods include both conventional (e.g., non-linear registration template construction approaches) and, more recently, deep-learning techniques, with deep-learning methods being initially established to focus on conditional factors, such as age; although, other joint factors have also been contemplated, such as gender and disease state [14]. Conventional methods produce templates using iterative processes [3], with templates typically constructed at a population level or for population-matched groups that reflect particular conditions, such as age. Using these methods, an individual template is constructed for each subgroup of interest (e.g., 80–90 year old cognitively normal participants). As such, these methods require huge amounts of data to produce templates that reflect a wide range or combination of clinical and demographic variables. Accordingly, reliance on pre-prepared publicly available population-based templates is widespread. However, it has become evident that population-based templates such as the Montreal Neuroimaging Institute (**MNI-152**) template do not accurately reflect global and regional brain structures among various sub- populations due to sampling biases inherent in the reference data [19]. Prior efforts using conventional methods to produce templates that reflect conditions such as age or disease state — for instance, the Mayo Clinic Adult Lifespan Template and Atlases [20] — have still suffered from significant limitations as they are often only produced for large age groupings (e.g., every 10 years) or for a particular status (e.g., patients with Alzheimer’s disease).

In contrast, deep-learning techniques for template construction offer a viable pathway to produce population level and conditional templates in a flexible manner. This could allow for the creation of templates that are conditioned on various combinations of parameters, such as age together with other demographic and clinical variables (e.g., biomarkers of disease progression), that are able to be jointly learned using a single model. While current methods offer a framework that can be conditioned on combinations of parameters, they currently lack the ability to geometrically link conditional templates to a common reference template or standard atlases.

This work evaluates key deep-learning methods that have recently emerged for conditional template construction [3, 14, 21], along with a method that builds on these key works, to establish how well they are able to represent underlying structural changes in the brain when conditioned on age as a demographic variable of interest. More complex sets of conditioning parameters (e.g., age, years of education and disease state) are of interest to neuroimaging analyses, although in the current study we focused on age to provide an evaluation of fundamental deep-learning conditional template construction methods.

### 1.1 Existing deep-learning methods

Dalca and colleagues have leveraged recent advances in deep-learning registration techniques [22] to produce a template construction method that induces templates by estimating network parameters that optimise a registration between templates and participant scans [3]. Their method is a two-stage network with: (i) the first-stage estimating a conditioned voxelwise intensity difference map that is used to adjust an average T1-weighted (**T1w**) intensity scan to produce conditional templates; and (ii) the second stage registering conditional templates to T1w participant scans to optimise network parameters. This enables the method to directly learn a function from the data that can jointly optimise templates tailored to specific conditions, such as age.

The method of Dalca *et al*. employs a diffeormorphic deformation framework in the second network stage to produce deformation fields that are smooth and invertible, thus creating registrations that promote topological consistency and preserve biological plausibility. To optimise the network, a global reconstruction loss term is utilised that calculates the mean squared error (**MSE**) between predicted and ground truth participant scans. In addition, various regularisation techniques are applied to the network that encourage deformations to be smooth, minimal, and centred, in order to promote consistency between conditional templates and participant scans. Firstly, a penalty term is applied to the average total magnitude of deformations to encourage deformations to be minimal. Secondly, a penalty term is applied to the first-order spatial derivatives of deformations to support overall spatial smoothness. Finally, a penalty term is applied to voxel-wise mean deformations to promote minimal average deformations across the dataset, which encourages the production of unbiased conditional templates that are centred in the distribution of deformations from templates to participant scans. This mean deformation constraint is designed to act globally across the dataset, where all training scans contribute to a single mean deformation field that is tracked and used to penalise the network.

Yu and colleagues developed a method to produce conditional probabilistic templates based on a single stage deformation network that deforms a group probabilistic template to condition specific templates [14]. This network uses ground truth segmentations of individual participant scans that have been modified to create one-hot encoded channel-wise segmentations. The ground truth channel-wise segmentations are then used to calculate similarities between individual participant scan segmentations and conditional probabilistic templates. Specifically, the loss function was formulated as the Soft-Dice coefficient [23], calculated between condition-specific probabilistic templates and one-hot encoded segmentations of individual participant scans. Similar to the method of Dalca *et al*., the method utilised a diffeomorphic deformation framework that estimates deformation parameters by scaling and squaring estimated stationary velocity fields. However, the method of Yu *et al*. does not utilise similar regularisation constraints to the method of Dalca *et al*. in order to encourage smooth or minimal deformations, with the loss function being comprised solely of the Soft-Dice coefficient. While this method has various applications, e.g., in conditional shape analyses of various brain regions, additional work is required in order to enable this method to effectively produce conditional structural templates (i.e., T1w conditional template images).

Dey and colleagues [21] extended the method of Dalca *et al*. by integrating adversarial training into a conditional template generation framework. In their method, a template is generated from a learned parameter array via a convolutional decoder. For conditional template synthesis, a covariate vector *z* (e.g., age) was incorporated using feature-wise linear modulation (**FiLM**). Specifically, given a feature map 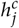 from the *j*th layer and channel *c*, the FiLM operation is defined as:

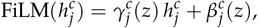

where 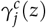 and 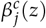 are scale and shift parameters computed from *z*. The generated template is then aligned to a target image through a registration module, in a similar fashion to the method of Dalca *et al*., yielding a diffeomorphic deformation field *ϕ* that warps the template to ground-truth participant scans. The registration module utilises a normalised cross-correlation (**NCC**) objective. In addition, regularisation terms are imposed on *ϕ* to encourage smooth, minimal, and centred deformations. To improve anatomical plausibility, the method employs an adversarial module — a PatchGAN-style discriminator [24, 25], conditioned on *z*, that differentiates between conditional templates registered to groundtruth participant scans and random ground-truth participant scans. The network is trained using a least-squares GAN (**LSGAN**) objective [26].

The aim of the current study was to extend these methods to produce conditional deformable structural T1w templates that are geometrically linked to both a group (unconditional) template and individual participant scans.

## 2 Methods

### 2.1 Data

A key consideration when preparing conditional templates using deep-learning methods is the study population used as it directly impacts the generalisability of methods and their effectiveness at accurately representing a target demographic. We focused our study on the production of conditional templates for an ageing cohort, given the applicability of this cohort to the analysis and prognosis of neurodegenerative conditions. We have utilised a cognitively normal (**CN**) subset of the Alzheimer’s Disease Neuroimaging Initiative (**ADNI**) study [27, 28] comprising 1534 T1-weighted MRI scans across 515 participants (55.02% female, 44.98% male; 50.5–95.7 years of age at scan date). The training dataset is comprised of 1230 scans, with the validation and testing datasets comprising 152 and 152 scans, respectively. In order to prevent cross-contamination among datasets, each of the datasets was constructed by first partitioning participants into each of the datasets such that any single participant was only represented in one of the training, validation or test datasets.

Data used in the preparation of this article were obtained from the Alzheimer’s Disease Neuroimaging Initiative (ADNI) database (adni.loni.usc.edu). The ADNI was launched in 2003 as a public-private partnership, led by Principal Investigator Michael W. Weiner, MD. The primary goal of ADNI has been to test whether serial magnetic resonance imaging (MRI), positron emission tomography (PET), other biological markers, and clinical and neuropsychological assessment can be combined to measure the progression of mild cognitive impairment (MCI) and early Alzheimer’s disease (AD). For up-to-date information, see www.adni-info.org.

#### 2.1.1 Dataset preprocessing

Our preprocessing pipeline included the following steps to address common MRI artefacts and enhance the comparability of scans across individuals:

1. ***Field of view adjustment*** (***FOV***) to remove lower head and neck voxels.
2. ***Skullstripping*** to remove non-brain voxels.
3. ***Bias-field adjustments*** to remove any bias-field artefacts remaining from acquisition.
4. ***Affine registration*** to a common space.

To focus analysis on the brain, we removed irrelevant sections of MRI scans, including the neck and lower portions of the head, using the *robustfov* tool from FSL [29]. We have then performed skullstripping with Freesurfer’s *SynthStrip* tool [30] to further remove non-brain voxels from scans.

The common MRI artefact of bias-field is addressed using FSL’s *FAST* tool [29], which is designed to efficiently segment tissue types at the same time as removing within-tissue low-spatial frequency intensity modulations. This has been well validated for the removal of bias-field artefacts.

Lastly, linear registration is applied to align all MRI scans to a common reference space, to standardise the global orientation and size, which is helpful for many neuroimaging analyses. We have utilised FSL’s *FLIRT* tool with 12 DOF for this task [31, 29].

### 2.2 Conditional template construction

We developed a purely geometric deep-learning approach to conditional template construction by relying on a two-stage deformation process, combining elements of the approaches presented in [3] and [14], whereby deformations are used to: (i) deform a group T1w template to create conditional template T1w volumes, and (ii) deform conditional templates to participant scans using a deep-learning registration approach. Figure 1 provides a schematic description of our template construction method: here, we have provided a simplified network diagram as well as a more detailed network schematic. A key advantage of our approach is the use of deformations in both network stages, so that conditional templates aim to be diffeomorphic to both the group T1w template and participant scans. Given that diffeomorphisms are invertible and differentiable, a network designed in this way should preserve the structural topology of the group template as it is deformed initially into conditional templates, and then subsequently to the participant scans.

**Figure 1:**
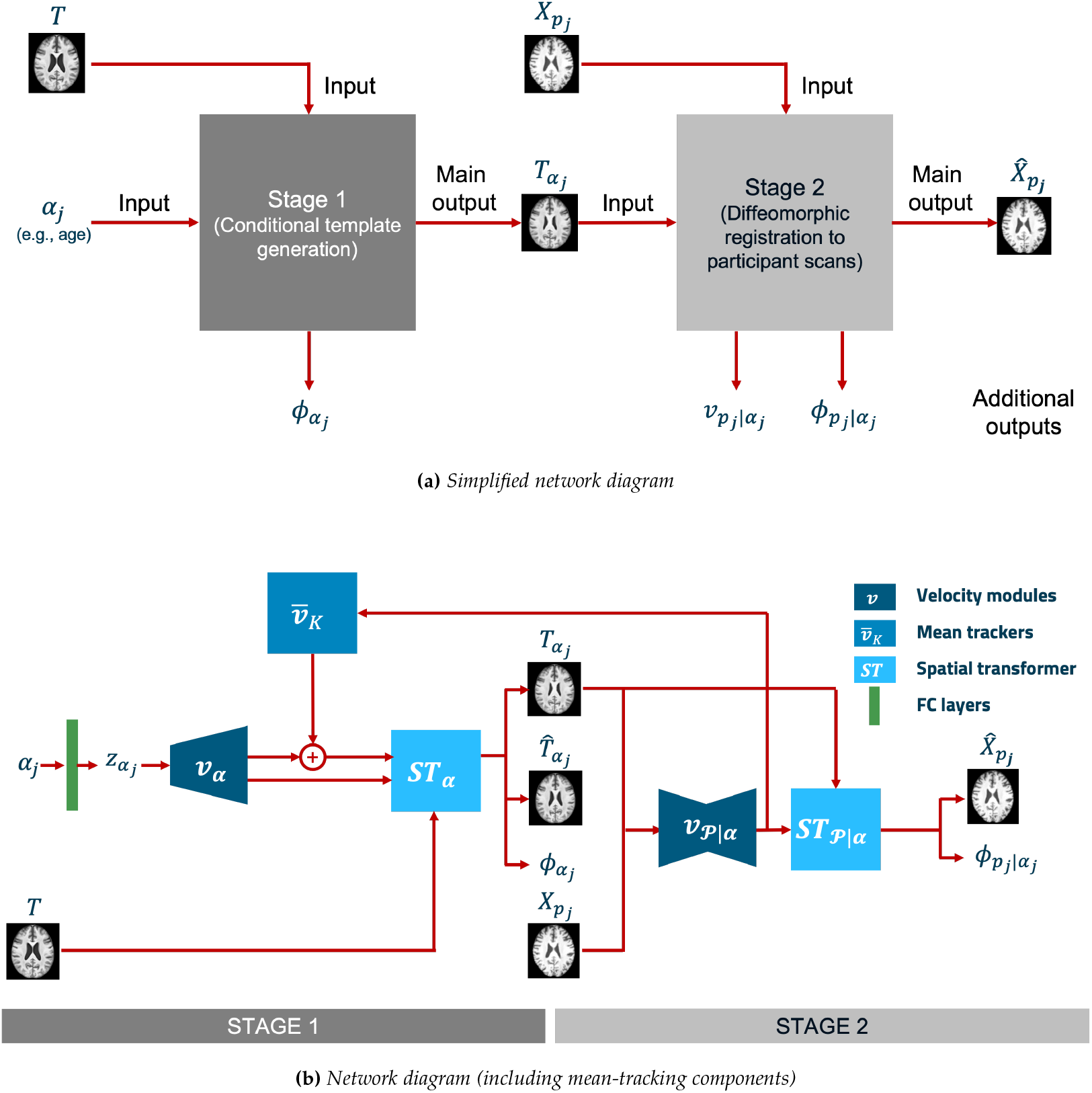
Network schematic for our two stage conditional template construction network. **Stage 1**: conditional parameters are passed to fully-connected (FC) layers to parameterise a latent distribution 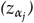, which is used to predict a conditional stationary velocity field,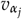. A spatial transformer layer integrates the conditional velocity field to produce a conditional deformation field, 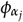 and deform the group template, T, to produce a conditional template, 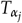 **Stage 2**: stationary velocity fields are predicted using a U-net architecture that ingests an individual participant scan, 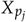, and conditional template, 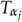, to produce a second stationary velocity field,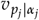. A second spatial transformer layer is then used to integrate 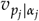 to produce a deformation field, 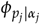 , and deform the conditional template, 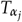 , to predict a participant scan,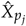. **Mean-tracking**: in addition to the above, velocity fields from stage 2 of the network are tracked by mean-tracking modules (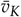in above schematic) in accordance with equation 5. Tracked mean velocity fields, 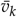, are combined with predicted stage 1 velocity fields in accordance with equation 6, which are used to predict a mean-adjusted conditional template, 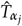 .

By contrast, the methods of Dalca *et al*. and Dey *et al*. apply a difference map to a linear average of *n* input scans, where *n* is set to 100 in their experiments. While this allows their methods to model non-geometric variability of conditional templates as a function of covariates, it does mean that these models are not geometrically constrained in the production of conditional templates, as boundaries can be effectively shifted in space via intensity changes.

#### 2.2.1 Group template

Given that our method is a purely geometric method, we require our group template to have sufficient anatomical detail to ensure that such details are not lost when generating conditional templates only through diffeomorphisms.

Accordingly, we require a group template that is generated using non-linear registrations. For the purposes of this work, we have used a subset of the training scans (20 female and 20 male), and produced the group template using the Advanced Normalization Tools (**ANTs**) [32] multivariate template construction algorithm, using a diffeomorphic symmetric normalisation registration (**SyN**) approach [33].

#### 2.2.2 Formulation of deformations

Our method builds upon the work of [3] and [14], but differed in that we utilised a single group T1w template, *T*, a discrete image with dimensions *D*x*H*x*W*, and applied deformations at both network stages to produce: (i) conditional templates (in stage 1), *T*_***α***_, for conditioning parameters *α*_*j*_ (e.g., age); and (ii) a deformed version of the conditional template to match the T1w scans, 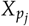 (in stage 2), associated with the *j*^th^ participant *p*_*j*_. Deformation fields, *ϕ*_***α***_ and 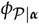 represent the mapping between: (i) a group T1w template and conditional templates, and (ii) conditional templates and participant scans, respectively.

The warping of the templates is defined as follows:

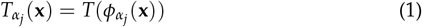

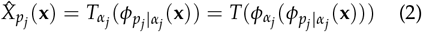

where interpolation is used to estimate values for non-integer coordinates.

We followed [34, 35, 36] to construct deformations from stationary velocity fields, *v*_***α***_ and *v* _ρ| ***α***_. Here, the deformation field is considered to be the solution of the differential equation:

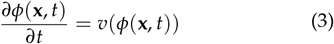

where *t* is time and *ϕ*(**x**, 0) = **x** defines the initial conditions. Deformation *ϕ*(**x**, 1) is calculated, in practice, by approximating the integration over the unit time interval with a set of scaling and squaring operations using the stationary velocity field vectors [37, 38, 34]. We use the simplified notation *ϕ*(**x**) or *ϕ* to denote *ϕ*(**x**, 1) in this paper.

#### 2.2.3 Model estimation

In order to estimate the deformation fields, *ϕ*_***α***_ and *ϕ*_P|***α***_, we followed a similar approach to [3], using a maximum likelihood estimation of stationary velocity field parameters, *v*_***α***_ and *v*_ρ|***α***_, as follows:

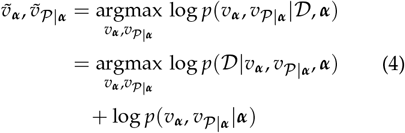

where the first term is the likelihood for the data, and assume that differences between the optimally warped conditional templates and participant scans have a Gaussian distribution; while the second term represents a prior (typically relating to smoothness) on the velocity fields, and 𝒟 is the dataset.

We modelled priors, *p*(*v*_***α***_, *v* _ρ|***α***_ |***α***), similarly to [3], under the assumption that deformations should be smooth, to maintain biological plausibility. In addition, the predicted conditional templates are likely to be geometrically biased towards the group template, which we want to avoid. Hence, we included terms to penalise: (i) deformation magnitude, (ii) deformation bending energy, and (iii) non-zero average deformations from conditional templates to participant scans that are clustered around a particular conditional parameter, such as age. Deformation magnitude and bending energy are calculated for both deformation fields, *ϕ*_***α***_ and *ϕ* _ρ|***α***_, while non-zero average deformations (from conditional templates to participant scans) are corrected for by producing a second conditional template that corrects for the average component of locally tracked stage 2 deformations (i.e., tracked across instances of similar age). Differences between the two conditional template images are used to encourage stage 1 deformation fields to be centered with respect to local stage 2 deformations.

While we modelled deformation magnitude equivalently to [3], there was a difference in the penalty terms that we used to encourage smoothness and minimal average deformations from conditional templates to participant scans.

#### 2.2.4 Model priors: smoothness

Smoothness was modelled by penalising bending energy as opposed to the first order spatial gradient. This is achieved by minimising the mean, across the image, of the squared Laplacian of the deformation field: (Δ*ϕ*)^2^.

The bending-energy formulation is based on the physics of thin elastic sheets or plates, where the optimisation of this energy minimises deformation curvature rather than penalising first-order spatial gradients. As a consequence, this more directly encourages smoothness, without penalising global changes in the scale or size of brain regions that may be biologically valid representations for various conditional templates or participant scans, while still controlling local deformations in order to preserve anatomical fidelity.

#### 2.2.5 Model priors: minimal average deformation from conditional templates to participant scans

Each conditional brain template should represent the average anatomy of all participants with the same conditioning parameter. To maintain the average position and shape of anatomical structures, the deformations from a conditional template to all corresponding participant scans must have a zero mean.

In order to achieve this we extended the idea of using global average deformation penalties to local condition-specific penalties. Accordingly, it is necessary to track average deformations from conditional templates to participant scans, at all locations, for all participants with a similar value of the conditional parameter, *α*_*j*_. This poses certain challenges: in order to penalise the network using local average deformation parameters for a range of conditional parameter settings, we need to store and track multiple instances of average deformation fields. For example, if age is the conditioning parameter, we need to track a separate average deformation field for a number of age bins, say each 5 year interval between 55 to 85 years of age. While this is feasible when the conditioning parameter is a simple scalar (as with age), when the dimensionality grows, the number of average deformation fields that need to be tracked will grow combinatorially. Additionally, when working with 3D images, batch sizes are typically too small to give a useful average within a batch, and so we need to update average deformation parameters across batches.

Despite these difficulties, we implemented this approach here by using soft binning for the conditional parameter space, ***α***, with one average deformation stored per bin. Updates to the averages were apportioned to the nearest two bins (nearest neighbours) using a linear weighting scheme, similar to approaches used for kernel density estimation.

Specifically, for this work we divided a one-dimensional conditional parameter space (age), ***α***, into a number of bins, where bin *k* stores an average deformation, 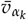(with average deformations tracked in velocity space). Equations 5–6 describe this simple case.

A soft binning scheme is used such that a weighting factor is defined that takes into account how close the age *α*_*j*_ is to the bin centre for bin *k*. This weighting factor, *ζ*_*k*_ (*α*_*j*_), is defined so that it is equal to 1 when *α*_*j*_ is equal to the *k*th bin centre, and falls off linearly to 0 as *α*_*j*_ approaches the neighbouring bin centres (for bins *k* − 1 or *k* + 1), and is 0 if it is further away.

Average deformations were iteratively tracked during each forward pass of the network, with average velocity estimates 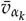 updated according to:

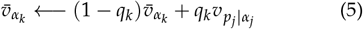

where the hyperparameter *q*_*k*_ combines a global value *λ*, which controls the update weighting (0 *< λ <* 1, and empirically set to 0.1), along with the soft binning weight, *ζ*_*k*_ (*α*_*j*_). That is, *q*_*k*_ = *λζ*_*k*_ (*α*_*j*_).

During each forward pass of the network, we construct centred conditional templates, 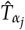, by combining the stage 1 velocity field, 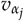, with the tracked average deformation velocity fields from stage 2 of our network as follows:

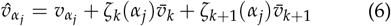

where *k* is chosen such that *α*_*j*_ lies between the centres of bins *k* and *k* + 1.

Deformation fields, 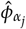, were derived (using equation 3), and applied to generate *mean-adjusted conditional templates*: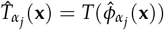. Following this, a penalty term was applied to 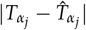 to encourage the network to predict velocities, and hence deformations and conditional templates, that are centered for conditioning parameters *α*_*j*_.

An alternative option for mean centring would be to penalise the average deformation parameters directly, however, we found empirically that this technique is less effective than using average deformation parameters to directly construct a centred conditional template, 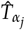. We then penalised differences between predicted and centred conditional templates, 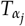 and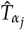, respectively.

#### 2.2.6 Conditional template network

Our method utilised a deep-learning probabilistic generative network in stage 1 to predict conditional stationary velocity fields, *v*_***α***_. We used fully connected layers to parameterise latent distributions, *p*(*z*|***α***), and then sample instances of *z*_***α***_. Convolutional transpose layers were then applied to produce predicted stationary velocity fields, *v*_***α***_. Following equation 6, velocity fields, *v*_***α***_, are then combined with tracked stage 2 velocity fields to produce 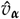 We used a spatial transformer layer [39] that: (i) integrates velocity fields, *v*_***α***_ and 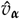, to produce deformation fields, *ϕ*_***α***_ and 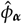; and (ii) deforms our group template to produce conditional templates, *T*_***α***_ and 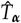. Following [22], we utilised a U-Net [40] that ingests conditional templates, *T*_***α***_, and participant scans, *X*_P_ , to predict stationary velocity fields, *v*_ρ|***α***_ A second spatial transformer layer was then used to integrate velocity fields, *v*_ρ|***α***_, to produce deformation fields, *ϕ*_ρ|***α***_, which were used to deform conditional templates, *T*_***α***_, to predict participant scans,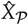.

Our network was constructed using the PyTorch deep-learning framework [41]. In order to optimise our network, we used the decoupled weight decay regularisation (AdamW) algorithm [42], with an initial learning rate of 10^−4^, *β*_1_ of 0.9, *β*_2_ of 0.999, and weight decay of 10^−2^. Our loss function consisted of 6 components: a reconstruction loss that calculates the reconstruction error (MSE) between predicted participant scans and ground truth participant scans (ℒ _recon_), the bending energy of deformation fields predicted for stage 1 and 2 of our network (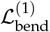 and 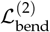, respectively), the global mean deformation magnitude predicted for stage 1 and 2 of our network (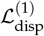 and 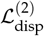, respectively), and reconstruction loss (MSE) between predicted conditional templates, 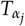 and mean-adjusted conditional templates,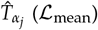. We used the Kornia Filters package [43] to calculate spatial gradients for 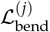. Deformation magnitude 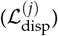 was calculated as the mean of the squared voxel-wise deformation magnitude (as used in [3]). Our full loss function was formulated as follows:

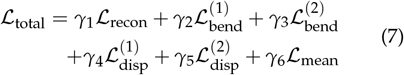

where *γ*_*j*_ are weightings applied to each component of ℒ _total_. We have set each of the weightings empirically, as follows: *γ*_1_: 1.0, *γ*_2_: 0.01, *γ*_3_: 0.01, *γ*_4_: 0.01, *γ*_5_: 0.01, and *γ*_6_: 1.0.

### 2.3 Evaluation

Important attributes of conditional templates include:

- topological consistency;
- structural fidelity;
- downstream utility; and
- accurate volumetric measurements for regions of interest.

We have evaluated these attributes, where reasonable to do so, for our method as well as for the methods of Dalca *et al*., Yu *et al*. and Dey *et al*. We have focused the evaluation on these methods when trained on the full dataset, described in section 2.1, using age as a conditioning parameter, although we have also evaluated each of these methods using both age and sex as conditioning parameters jointly, as well as female-only and male-only partitions of the full dataset. In addition, we have evaluated our method using 25% and 50% of the training dataset (308 and 616 training scans, respectively), as well as with training data that was corrupted by: (i) random noise to target a signal to noise ratio (**SNR**) that is 30–40% that of the raw scans, and (ii) making random contrast adjustments to the raw scans by raising an input scan’s voxel intensities to *γ*; i.e. 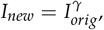 where *γ* = *e*^*β*^ and *β* ∼ 𝒰 ( − *a, a*) where *a* is set to 0.3. We have also evaluated the impact of each of the loss function components (excluding ℒ_recon_) by setting the weighting parameters, *γ*_2_–*γ*_6_, to zero iteratively, such that for each experiment, one of *γ*_2_–*γ*_6_ is equal to zero. These results are presented in the supplementary materials.

#### 2.3.1 Topological consistency

The preservation of topology promotes anatomical accuracy and biological plausibility, as adult brains that are free from obvious pathologies share the same essential topology, despite substantial geometric heterogeneity between individual brains. Conditional templates that maintain the brain’s topology are more likely to represent plausible anatomical variations, as this should be preserved as conditioning parameters such as age vary.

To measure the topological consistency between different brain representations, the determinant of voxel-wise Jacobian matrices, *J*_*ϕ*_(*j*) =∇ _*ϕ*_(*j*) ∈ ℛ ^3×3^, was calculated for deformation fields between images (e.g., unconditional and conditional templates). The determinant, |*J*_*ϕ*_(*j*)| , captures the local change in volume, such that deformations that do not preserve local topology have |*J*_*ϕ*_(*j*) | ≤0. In addition, we show the distribution of the Jacobian determinants across the images to assess the scale of volume changes that each method was able to produce on a voxelwise basis.

#### 2.3.2 Structural similarity

Well-constructed conditional templates should have good structural fidelity and be similar to individual brains (e.g., the shape and intensity of corresponding brain structures should be similar). Conditional templates with similar (adjacent) conditional parameters should exhibit greater structural similarity than those where the conditional parameters differ more.

The structural similarity of adjacent conditional templates can be measured using the Structural Similarity Index Measure (**SSIM**) [44], which extracts and compares three key images features: (i) luminance, *l*(· ); (ii) contrast, *c*(·); and (iii) structure, *s*(· ). It then combines these to produce a single overall measure: *f* (*l*(**x, y**), *c*(**x, y**), *s*(**x, y**)) for images **x** and **y** (more details can be found in supplementary materials section S1).

While there should be substantial structural similarity between adjacent templates, these should not be perfectly similar, as some differences are to be expected (e.g., due to age). There should exist a balance between structural similarity being consistently high between adjacent templates and changes of interest being apparent. We report SSIM before and after non-linear registration is applied, as this can provide an assessment that is less influenced by welldefined geometric changes in brain structure that can be captured and removed by the registration. The SyN algorithm was used to perform the registrations, with default hyperparameters as provided by the ANTs implementation.

The root mean squared difference (**RMSD**) between voxel intensities is also reported, as it is a straightforward measure (calculated equivalently to RMSE) that is a purely local, intensity-driven similarity measure. RMSD is an effective tool for detecting even small differences in the anatomy or appearance between conditional templates.

In addition, we have reported average Dice Similarity Coefficient (**DSC**) results, calculated for regions of the Deiskan Killiany (**DK**) atlas, to assess the degree to which methods are able to preserve and recover regional brain structure. In order to measure the DSC for each method, we have non-linearly registered the DK T1 template to each conditional template, and used the resulting deformations to move the DK atlas labels to each conditional template space. Conditional template segmentations are computed using the FastSurfer segmentation tool [45]. We have performed visual quality control (**QC**) of all the segmentations generated by the FastSurfer tool for this and other evaluations in this work. Two failures were detected and removed as a result of this process.

#### 2.3.3 Downstream utility

We have used conditional templates as part of a downstream binary classification pipeline to discriminate CN versus MCI/AD participants. We have used a cohort of 675 participants from ADNI3 (56.4% CN, 32.3% MCI, 11.3% AD). To assess the utility of features derived from conditional templates for the purposes of classification, we have used Jacobian determinant maps derived from non-linear registration deformation fields (taking a similar approach to Hua and colleagues [46]), with the four following options: (i) direct registrations from individual participants to our unconditional group template (**direct**); (ii) chained registrations from individual participants to an appropriate age-related conditional template, and then to our unconditional group template (**chained**); (iii) registrations from individual participants to an appropriate age-related conditional template, with Jacobian determinant maps moved to our unconditional group template space (**age**); and (iv) a concatenation of the features derived from (ii) and (iii) (**concatenated**).

To distil the features derived from spatial maps of Jacobian determinants, we have created feature masks by performing a voxel-wise ANOVA test on Jacobian determinant maps (attached to our unconditional group template space) using an independent hold-out sample of 80 participants (40 CN, 25 MCI, and 15 AD), with an uncorrected p-value threshold of 0.001. This hold-out sample was selected to provide a balanced representation of the clinical groups and was reserved solely for feature selection. The remaining 595 participants have been utilised in our classification pipeline.

For each feature set, we have adopted a leave-one-out (**LOO**) strategy, training linear kernel SVMs (CN vs. MCI/AD) for each fold. We have performed a one-sided binomial test to evaluate whether there is an improvement in the classification accuracy (i.e., proportion of correct outputs) using the conditional template options (i.e., chained, age, and concatenated) versus the direct option.

#### 2.3.4 Volumetric measurements for regions of interest

Volumes of various regions of interest (**ROIs**) are of considerable interest in understanding developmental and degenerative processes. Therefore, the accuracy and reliability of these volume measures demonstrates whether conditional templates are practically effective at capturing anatomical change as a function of the conditioning parameter(s), such as age. Such validation is important for the adoption of methods in both research and clinical settings.

Volumes for the conditional template ROIs were obtained by applying a segmentation method independently to each template. We also normalised all volumes of a particular ROI by the mean volume of that ROI across all conditional templates, independently for each method. This provides a simpler comparison across methods and different ROIs (i.e., to show proportional changes in average volume), without undue influence from the values at the ends of the conditional parameter range, which are often under-represented and hence poorly estimated.

More specifically, the normalised volume is formulated as follows:

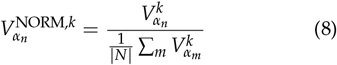

where *k* is an index representing the ROI, and *n, m* ∈ *N* are indices representing conditional parameters (e.g., age).

Ground truth volumes were also obtained by segmenting the individual training scans. These were grouped into bins, based on conditional parameters values (age), and the average values from each bin were then normalised using equation 8. More specifically, the average normalised volume for training scans in each bin is:

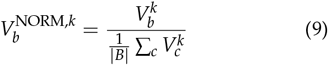

where *b, c* ∈ *B* represents the set of bins partitioning the training scans and 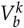 is the average volume for ROI_*k*_ in bin *b*.

These training scans volumes act as a simple unbiased estimate of what the volume should be at each age, and are plotted along with the corresponding standard deviation. This provides a standard for assessing the volumes obtained from each of the methods, to see if they are able to accurately capture morphological variations in brain anatomy.

## 3 Results

Evaluation criteria, as described in section 2.2, were assessed for our method as well as for the methods of Dalca *et al*. [3], Yu *et al*. [14] and Dey *et al*. [21]. For the method of Dalca *et al*., we implemented their conditional template network using the PyTorch deeplearning framework, following [3] and code available from their website^1^. For the method of Yu *et al*., we utilised the code available from github^2^, with adjustments made to their codebase to accommodate our dataset (see section 2.1). For the method of Dey *et al*., we utilised the code available from github^3^, and again made adjustments for our dataset.

### 3.1 Topological consistency

Jacobian determinants of deformation fields from unconditional to conditional templates may only be computed for our method and the method of Yu *et al*. The methods of Dalca *et al*. and Dey *et al*. do not use a deformation field to produce conditional templates, as they only use deformation fields to deform conditional templates to participant scans. However, we have also computed Jacobian determinants of deformation fields for conditional templates to predicted participant scans for our method and the methods of Dalca *et al*. and Dey *et al*., given that the method of Yu *et al*. is a one-stage deformation network.

Figure 2 displays distributions of the Jacobian determinants for the stage 1 conditional deformation fields (ours and Yu *et al*.) as well as stage 2 deformation fields (ours, Dalca *et al*. and Dey *et al*.). From these results, it is evident that the deformations produced by our method, in both cases, and for the method of Yu *et al*., are well centred around 1.0. Additionally, our method did not produce negative Jacobian determinants (range of values were 0.119 to 2.298 for stage 1 and 0.121 to 2.301 for stage 2), indicating that it can effectively maintain topological consistency through deformations. For stage 2, the method of Dalca *et al*. exhibited a much wider range of Jacobian determinants, with determinant values spanning -2.7 to 22.2. A number of voxels exhibited negative Jacobian determinants within these fields, with an average of 473 voxels for each of the stage 2 deformation fields in the test dataset. This indicates that topological inconsistencies are introduced between the conditional template and predicted participant scans, which could be a result of less regularised registrations or topological alterations in their conditional templates. Similarly, the method of Dey *et al*. also produced a wide range of Jacobian determinant values in stage 2, spanning -0.7 to 31.9, and while this method still produced negative Jacobian determinants, there were only 36 instances across the 152 participants in the test dataset. While the method of Yu *et al*. produced a distribution centred around 1.0, the Jacobian determinant values ranged from -0.2 to 3.4, with an average of 9 voxels for each of the conditional template deformation fields exhibiting negative Jacobian determinants.

**Figure 2:**
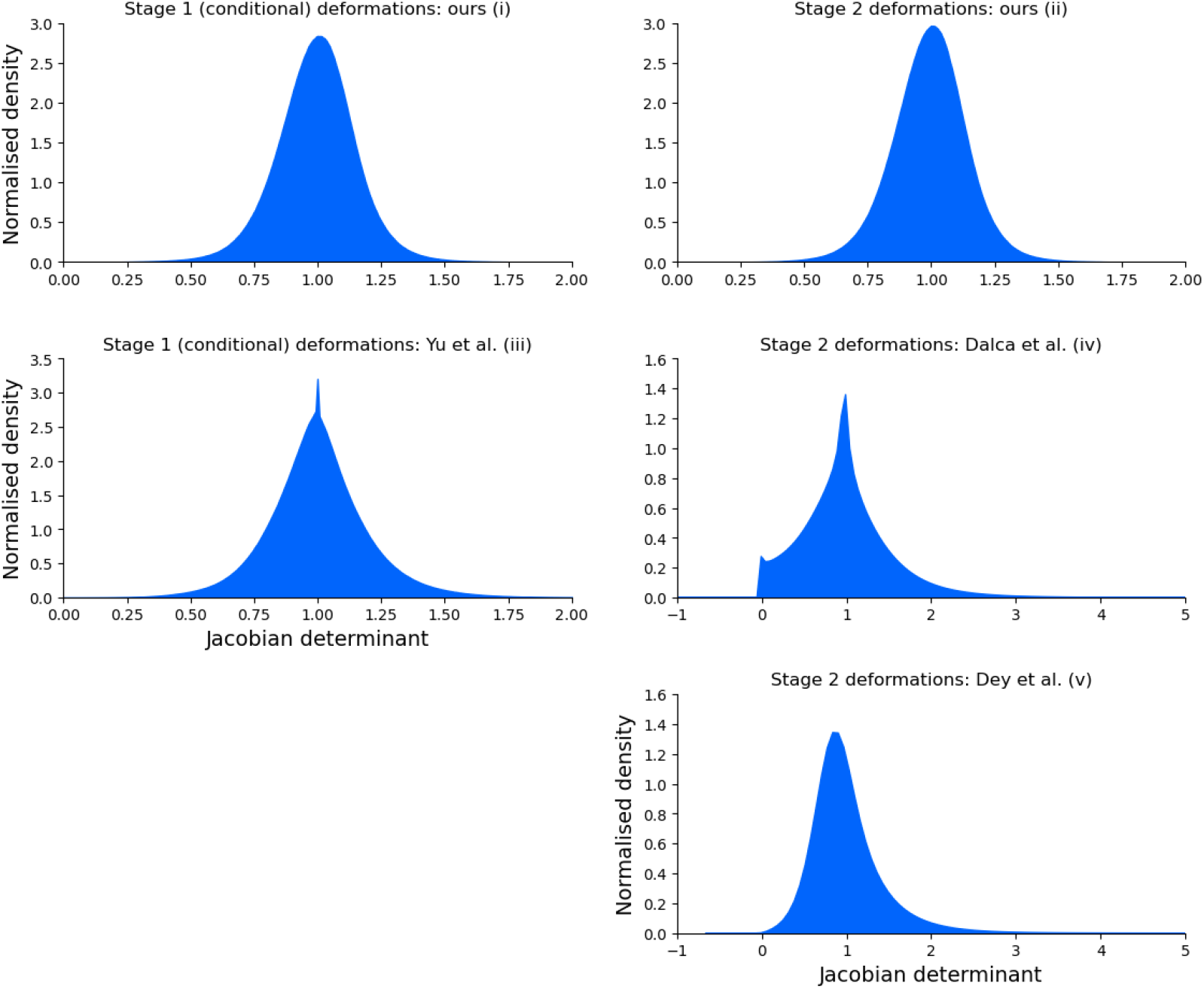
Distributions of Jacobian determinants for: (i) our method for deformation fields from unconditional to conditional templates, and (ii) conditional templates to predicted participant scans; (iii) deformations from unconditional to conditional templates for the method of Yu et al.; (iv) deformations from conditional templates to predicted participant scans for the method of Dalca et al.; and (v) deformations from conditional templates to predicted participant scans for the method of Dey et al.

Spatially, the method of Dalca *et al*. produced negative Jacobian determinants that were distributed predominantly within the cortical grey-matter when averaged across stage 2 deformation fields produced for the test dataset. For both the methods of Yu *et al*. and Dey *et al*., the negative Jacobian determinants produced for the conditional template deformation fields and stage 2 deformation fields, respectively, were sparsely distributed, and for the method of Yu *et al*., they were typically located in the posterior cortex. These results are available in section S14 of the supplementary materials.

Jacobian determinants were also calculated for each of these methods using both age and sex as conditioning parameters jointly, as well as for female- and male-only partitions of the training dataset. In addition, we have computed Jacobian determinants for our method using 25% and 50% of the training dataset, and when using training data that was corrupted with random Gaussian noise or with random contrast adjustments. Finally, Jacobian determinants were also calculated for our method where the loss function components, *γ*_2_–*γ*_6_, were iteratively set to zero, as described in section 2.3. The results for these experiments are broadly similar to those for the original, full training dataset, although the method of Yu *et al*. produced distributions of Jacobian determinants with lower variance than those produced using the full training dataset with age as the only covariate, with the distribution for the two covariate model (i.e., age and sex) and male-only data exhibiting much lower standard deviation when compared to the distribution for the full dataset with just age as a covariate (0.04 and 0.20, respectively). Additionally, when setting *γ*_3_ to zero for our method, while negative Jacobian determinants were not produced, the distribution of Jacobian determinants for both conditional template deformations and stage 2 deformations were skewed, spanning 0.03 to 14.67 and 0.04 to 13.22, respectively.

Figures showing these additional Jacobian determinant results are available in the supplementary materials as follows: section S2 for reduced and corrupted datasets, section S5 for models trained with age and sex as covariates, section S8 for sex-specific datasets, and section S11 for adjusted loss components.

### 3.2 Structural similarity

We calculated SSIM for our method, as well as the methods of Dalca *et al*. and Dey *et al*. The method of Yu *et al*. is not amenable to these comparisons as the conditional templates that are produced are conditional probabilistic atlases.

Figure 3 displays SSIM results for our method and the methods of Dalca *et al*. and Dey *et al*. for adjacent conditional templates separated by age gaps of 5 years. While SSIM metrics are above 0.95 for each method, on both pre- and post-registration bases, SSIM performance for the method of Dalca *et al*. changes very little with registration and degrades at the tails of the age distribution. Given that the method of Dalca *et al*. generates conditional templates from a difference map, the network is less constrained compared to producing conditional templates through purely geometric deformations of a group template. Despite this, the method of Dey *et al*., which also uses a difference map to generate conditional templates, was able to produce more consistent SSIM results across the age distribution.

**Figure 3:**
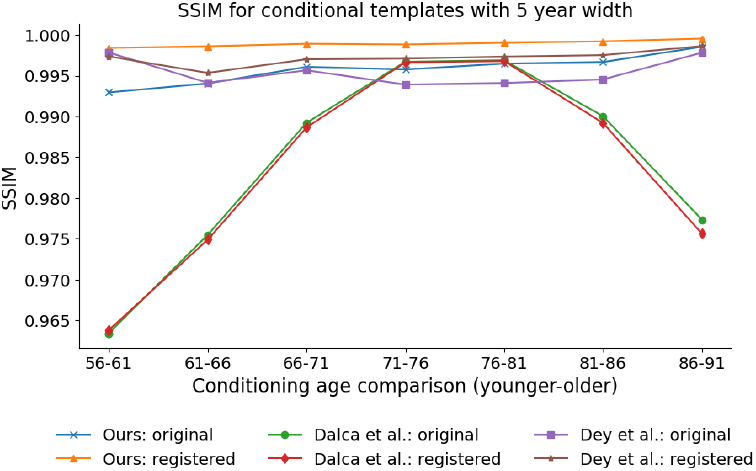
Structural Similarity Index Measure (**SSIM**) for predicted conditional templates separated by 5 years, on both a pre- (original) and post-registration basis. For the registered images, each lower age conditional template has been registered to the next higher age conditional template, with the registered image and the higher age template used to calculate SSIM.

Figure 4 displays RMSD results. It should be noted that the RMSD results are reasonably close to zero for each of the methods, which indicates that there is considerable similarity between each of the pairs of conditional templates. However, a low RMSD value may result from negligible differences in every voxel, or more likely, a strong difference in a small number of voxels (e.g., near the edges of various ROIs). Of interest, our method produced higher (less similar) RMSD results when calculated on a pre-registration basis. Such a result may be due to greater differences exhibited by macro-brain-structures (e.g., the lateral ventricles) as the conditioning age parameter is adjusted; which is supported by the post-registration RMSD results, where the registration can remove such geometric changes, being comparable to the RMSD results for the method of Dalca *et al*. While the method of Dey *et al*. did see an improvement in RMSD on a post-registration basis, it was lower than the improvement exhibited by our method, and is less evident at the ends of the age distribution, particularly at the lower-end.

**Figure 4:**
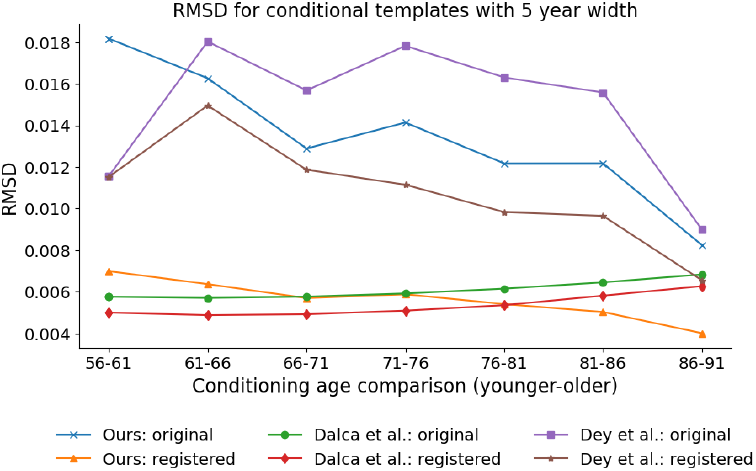
Root mean squared difference (**RMSD**) for predicted conditional templates separated by 5 years, on both a pre- (original) and post-registration basis. For the registered images, each lower age conditional template has been registered to the next higher age conditional template, with the registered image and the higher age template used to calculate RMSD.

For our method, voxel-wise absolute differences between adjacent conditional templates in conditional parameter space (where one conditional template is registered to the other) exhibit a spatial distribution where the greatest differences are typically expressed around large structures (e.g., the lateral ventricles). For the method of Dey *et al*., the spatial distribution of voxel-wise absolute differences demonstrate that the greatest differences are not only located around large structures like the lateral ventricles, but there are also significant differences distributed near sulci in the cortex. For the method of Dalca *et al*., the voxel-wise absolute differences exhibit a smaller range, although moderate values are more widely distributed. See illustrative figure in supplementary materials section S14.

We have also calculated SSIM and RMSD for each of these methods using both age and sex as conditioning parameters jointly, as well as for female- and male-only partitions of the full training dataset. In addition, we have computed SSIM and RMSD for our method using 25% and 50% of the training dataset, and when using training data that was corrupted with random Gaussian noise or with random intensity contrast adjustments. Finally, SSIM and RMSD were also calculated for our method where the loss function components, *γ*_2_–*γ*_6_, were iteratively set to zero, as described in section 2.3.

For models trained using age and sex as conditioning parameters jointly, the SSIM results for our method and the method of Dey *et al*., and the RMSD results for our method and the methods of Dalca *et al*. and Dey *et al*. are similar to those produced for models trained using only age as a conditioning parameter. The SSIM results for the method of Dalca *et al*., while very similar on a pre- and post-registration basis, do not exhibit a similar decline at the ends of the age distribution.

When using sex-specific, reduced or corrupted training datasets, the results for our method are similar to those produced using the full, original training dataset, for both SSIM and RMSD, although for the male-only data there was a decline in SSIM and RMSD performance at the lower end of the age distribution on a pre-registered basis.

When adjusting loss components *γ*_2_–*γ*_6_ iteratively to zero, both SSIM and RMSD performance remained mostly consistent, with two exceptions: (i) when setting *γ*_3_ to zero, there was a decline in both SSIM and RMSD performance on both a pre- and post-registration basis, and (ii) when setting *γ*_6_ to zero, there was a reduced difference between pre- and post-registration performance for both SSIM and RMSD, with performance for both metrics, similar to our method with all loss components active on a post-registration basis.

The SSIM results for the method of Dalca *et al*. exhibited a similar parabolic shape across the age distribution for both the female- and male-only data as those produced for the full training dataset, although the rate of decline in the SSIM metric was less pronounced at the ends of the age distribution, particularly for the male-only data. While the method of Dey *et al*. demonstrated improved SSIM and RMSD performance on a post-registration basis (compared to pre-registration performance) for both the female- and male-only data, the SSIM and RMSD results were generally lower than those produced for the full training dataset, particularly for the male-only data.

Figures showing the results for SSIM and RMSD in these various cases are available in the supplementary materials as follows: section S3 for reduced and corrupted datasets, section S6 for models trained with age and sex as covariates, section S9 for sexspecific datasets, and section S12 for adjusted loss components.

We have calculated the average DSC for our method, as well as the methods of Dalca *et al*. and Dey *et al*. The method of Yu *et al*. is similarly not amenable to these comparisons as the conditional templates that it produces are conditional probabilistic atlases.

Figure 5 displays the DSC results where DSC is averaged across conditional templates produced by a method for each integer age in the range 55–95 years, inclusive. These results demonstrate that while each of the methods are able to preserve anatomical fidelity reasonably well across certain structures (e.g., for the lateral ventricles, thalamus, and brainstem, DSC results were greater than 0.85 for all methods), they struggle with challenging or heterogeneous regions (e.g., cerebral cortex).

**Figure 5:**
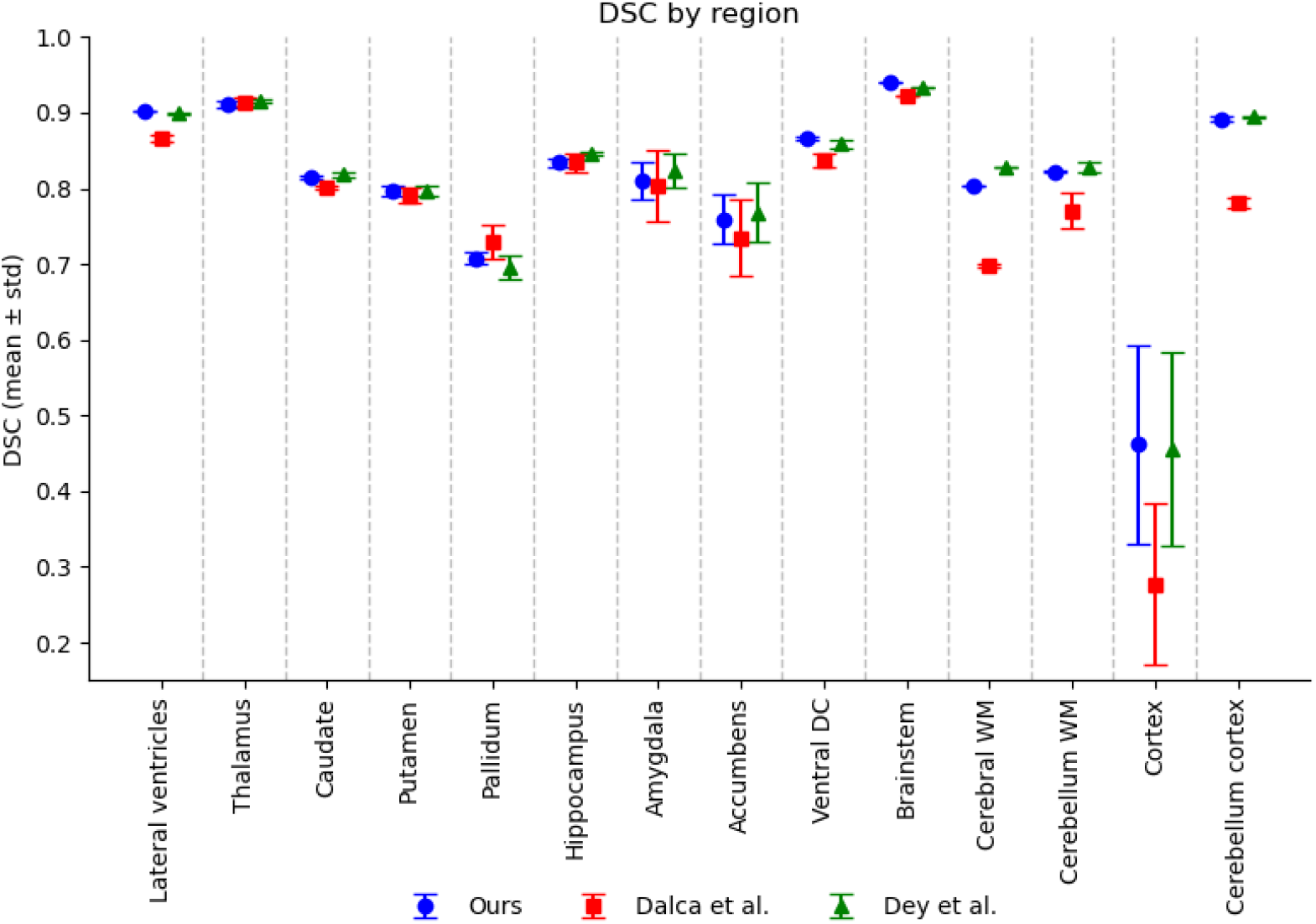
Average Dice Similarity Coefficient (**DSC**) results, calculated for regions (or merged regions) from the Deiskan Killiany (**DK**) atlas. In order to measure the DSC for each method, we have non-linearly registered the DK T1 template to each conditional template, and used the resulting deformations to move the DK altas labels to each conditional template space. Conditional template segmentations are computed using the FastSurfer segmentation tool.

We have also evaluated the utility of the conditional templates produced by our network for the task of registration by registering the group template and conditional templates produced by our network to 50 participant scans randomly selected from our test dataset, using the SyN algorithm [33]. We assessed image similarity by calculating the root mean squared error (**RMSE**), normalised cross-correlation (**NCC**) and mutual information (**MI**) between the participant scans and the registered templates. The distributions of the differences between these metrics for the conditional and group templates are shown in the supplementary materials section S15, and demonstrate that the conditional templates produced by our network are superior to an unconditional template for this task.

In addition, we have evaluated how well individual participants align to conditional templates across methods. We have non-linearly registered each participant scan in the test dataset to conditional templates (produced for the age of the participant at scan date using our method, as well as the methods of Dalca *et al*. and Dey *et al*.) using the SyN algorithm. We have then calculated RMSE, NCC and MI between each registered participant scan and its corresponding conditional template. Box plots for each of these metrics and methods are shown in the supplementary materials section S16.

Our results indicate that both our method and the method of Dey *et al*. outperform the method of Dalca *et al*. across each of RMSE, NCC, and MI. There is negligable difference observed between our method and the method of Dey *et al*. for RMSE, however, the method of Dey *et al*. is slightly better than our method for NCC and MI.

We note that for the method of Dey *et al*., their reconstruction loss was driven by an NCC criterion, while for our method and the method of Dalca *et al*., the reconstruction loss was driven by an MSE criterion. We have compared using NCC in place of MSE for our reconstruction loss, as well as FiLM conditioning in stage 1 of our network (as per the method of Dey *et al*.), however, we have found that these changes have not led to systematic improvements in conditional template generation.

### 3.3 Downstream utility

We have assessed the utility of features (i.e., Jacobian determinants) derived using conditional templates for a downstream binary classification task for our method, as well as the methods of Dalca *et al*. and Dey *et al*. The method of Yu *et al*. is not suitable for this analysis as the conditional templates that are produced are conditional probabilistic atlases.

Table 1 shows the accuracy and binomial test results for this classification task. Each of the conditional template options (i.e., chained, age, and concatenated) produced higher accuracy results than the direct option, with the concatenated features demonstrating a 4.2% improvement (p-value of 0.0002).

**Table 1:**
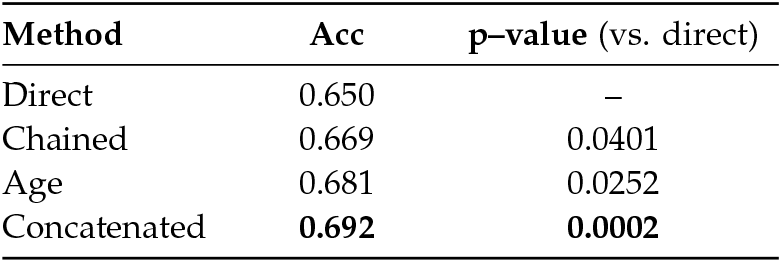
Classification accuracy and binomial test comparisons (CN vs. MCI/AD). The baseline direct option classification accuracy is shown in the first row, with the conditional template options (i.e., chained, age, and concatenated) recording classification accuracy and p-values from binomial tests (vs. the direct option).

### 3.4 Volumetric measurements for regions of interest

We focused our volumetric measurements on particular ROIs and tissues that are implicated in ageing and neurodegeneration, namely: (i) hippocampus, (ii) lateral ventricles, (iii) cortical grey-matter, and (ii) white-matter. These measurements provided a benchmark to compare with those reported in the literature [3, 14]. To measure structural volumes for each of the deep-learning methods, we adopted two approaches. For our method and the methods of Dalca *et al*. and Dey *et al*., we have used the FastSurfer segmentation tool to segment and calculate volumes for each conditional template produced by these deep-learning models as trained on the ADNI dataset (described above in section 2.1). For the method of Yu *et al*., we calculated volumes by thresholding the predicted conditional probabilistic atlases and summed voxels across each channel that reflected a particular ROI; we believe that this is the most direct way to calculate volumes for ROIs as the method outputs conditional probabilistic atlases that are deformed from an unconditional probabilistic atlas, and not intensity-based images suitable for input to FastSurfer. We also used FastSurfer to segment and calculate volumes directly from the training data. For each of the methods and training scans, we then normalised volumes by dividing each volume by the mean of all volumes, across all bins, independently for each ROI and method (see equations 8 and 9).

Figure 6 depicts normalised volumes, for each ROI, from each of the deep-learning methods being evaluated, as well as volumes calculated directly from the training dataset, using bins with a width of 2 years. These results demonstrate that each of the methods performed differently across the various ROIs considered. The methods of Yu *et al*., Dey *et al*., and our method produce volumes for the lateral ventricles as a function of age that are more consistent with those obtained from the training data as compared to the method of Dalca *et al*.. Our method and the method of Yu *et al*. performed slightly better than the method of Dey *et al*. at the lower end of the age distribution, while our method and the method of Dey *et al*. more closely follow the training data at the upper-end of the age distribution. For white-matter volumes, the method of Dalca *et al*. produced volume differences that are less consistent with the changes exhibited in the training data when compared to the methods of Yu *et al*., Dey *et al*., and our method. However, all of the methods display some underestimation of the changes compared to those measured directly from the training dataset scans. The methods of Dalca *et al*. and Dey *et al*. showed better agreement with the training dataset results for cortical grey-matter volume than both the method of Yu *et al*. and our method, although none of these methods show the same degree of change as those measured directly from the training dataset. For hippocampal volumes, our method produced very minimal changes, which are inconsistent with what we would expect to see as age increases. While both the methods of Yu *et al*. and Dalca *et al*. exhibited larger changes, they potentially underestimate the change shown by the volumes directly measured from the training images. However, with low numbers of participants in the higher and lower age bins, the values at the ends of the age range are not as trustworthy as those in the middle of the age range. Despite this, the method of Dey *et al*. was able to track hippocampal volumes of the training data remarkably well.

**Figure 6:**
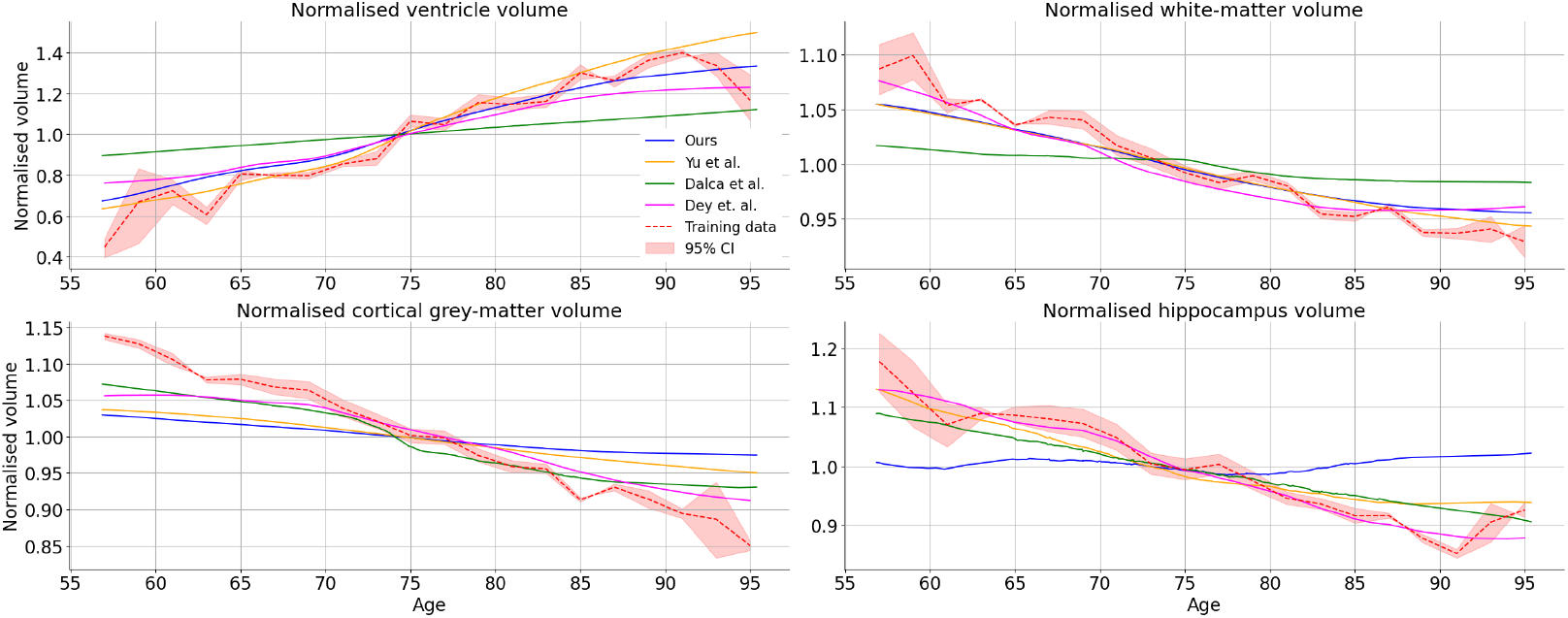
Volumetric measurements for various ROIs: cortical grey-matter, white-matter, lateral ventricles and hippocampus. Volumes are presented on a normalised basis, where volumes are normalised by the mean of all volumes across the range of conditional parameters, independently for each ROI and method. In addition, a 95% confidence interval is also presented for the empirical estimates made from training data within 2-year age bins.

Volumetric measurements were also made for each of these methods using both age and sex as conditioning parameters jointly, as well as for female- and male-only partitions of the training dataset. In addition, we have made volumetric measurements for our method using 25% and 50% of the training dataset, and when using training data that was corrupted with random noise or with random contrast adjustments. Finally, volumetric measurements were made using our method where the loss function components, *γ*_2_–*γ*_6_, were iteratively set to zero, as described in section 2.3.

While these results were similar to those produced using the full, original training dataset, with age as the only conditioning parameter, there were some exceptions. Firstly, for our method, while producing similar results for hippocampal volumes on the female-only data and for the model trained using age and sex as conditioning parameters (where age is varied while the female conditioning parameter is held constant), produced volumes for the male-only data, and for the model trained using age and sex as conditioning parameters (where age is varied while the male conditioning parameter is held constant), that erroneously increased across the age distribution. For our method, when setting *γ*_6_ to zero (i.e., excluding ℒ_mean_ from the loss function), we observed underestimation of normalised ventricle and white-matter volumes compared to these volumes when estimated by our method with all loss function components active.

Figures showing these additional volumetric results are available in the supplementary materials as follows: section S4 for reduced and corrupted datasets, section S7 for models trained with age and sex as covariates, section S10 for sex-specific datasets, and section S13 for adjusted loss components.

## 4 Discussion and conclusion

The use of deep-learning methods for the construction of conditional brain templates has many advantages. In order for such methods to gain wider acceptance and provide meaningful utility to both researchers and clinicians, our analysis suggested that they require further refinement. For instance, while methods may be able to capture some morphological differences associated with the ageing adult brain (e.g., volumetric changes in the ventricles), it is also apparent that they do not accurately represent all changes across the brain (e.g., volumetric changes in cortical grey-matter). Moreover, although our method produced visually high quality conditional T1w templates, these failed to capture important ageing characteristics of hippocampal degeneration. Conversely, while the methods of Dalca *et al*. and Yu *et al*. were able to more accurately capture the volumetric distribution of hippocampal volumes, these methods either do not produce conditional T1w templates (Yu *et al*.) or produce conditional T1w templates with lower image quality when compared to our method (as demonstrated by the results in figure 3).

When we consider topological consistency, our method outperformed both the methods of Dalca *et al*. and Dey *et al*. for deformations from conditional templates to individual participant scans (stage 2 of the respective networks) — for these deformations, our network did not produce any negative Jacobian determinants while the methods of Dalca *et al*. and Dey *et al*. produced a number of very small and negative Jacobian determinants, which indicates that there were several instances of topological inconsistency between conditional templates and individual scans, with the method of Dalca *et al*. producing an average of 473 voxels that exhibited negative Jacobian determinants for each of the predicted deformation fields. While the method of Dey *et al*. produced far fewer negative Jacobian determinants than the method of Dalca *et al*., there were still 36 instances across the test dataset, which indicates that this method is still prone to the introduction of topological inconsistencies. Additionally, while the distribution of Jacobian determinants for stage 2 deformations was reasonably symmetric for our method, both the methods of Dalca *et al*. and Dey *et al*. produced a right-skewed distribution with Jacobian determinant values spanning -2.7 to 22.2 and -0.7 to 31.9, respectively. For deformations from the group template to conditional templates (stage 1 of our network), while our method did not produce any negative Jacobian determinants, which indicates that it was able to maintain topological consistency between group and conditional templates, the method of Yu *et al*. produced an average of 9 voxels with negative Jacobian determinants for each of these deformation fields. Additionally, for our method, the distribution of Jacobian determinants is similar and symmetrical across both network stages, which may imply that it is effective at factoring out changes that are common across the population and associated with age, from those that are associated with individual anatomical differences.

From a structural similarity perspective, both our method and the method of Dey *et al*. can produce adjacent conditional templates that share substantial similarities as conditioning parameters are adjusted, with high SSIM metric values for adjacent conditional templates, prior to any additional registration, across the range of age comparisons considered. Furthermore, after additional non-linear registration, both our method and the method of Dey *et al*. produced SSIM metric values that were consistently improved compared to the pre-registration values, which we would see if there was useful and measurable geometric change being captured with varying age that registration could undo. Conversely, the method of Dalca *et al*. produced similar SSIM metric values both before and after registration. This implies that the age-related (conditional parameter) changes were not able to be easily explained in terms of geometric changes, such as the size and location of brain structures, by the registration process, even though we would expect to see such age-related changes in geometry. Additionally, SSIM results for the method of Dalca *et al*. exhibited decreasing similarity at both ends of the age distribution, which indicates that greater structural discrepancies were observed, between adjacent 5-year intervals, at the edges of the conditional parameter space.

The volumetric measurement results for several ROIs that are known to vary in the ageing brain are mixed, with each method having advantages and disadvantages over other methods across different structures. For instance, our method could track the distribution of volumes for structures like the ventricles with good accuracy, and to a lesser extent, for white-matter tissue. However, our method tracked the distribution of hippocampal volumes poorly, and was also deficient in tracking the distribution of cortical grey-matter tissue volume. On the basis of volumetric measurements, the methods of Dey *et al*. and Yu *et al*. were the strongest approaches when considering the various structures examined, with good accuracy, compared to the empirical ranges estimated from the training scans, for the volumes of the ventricles, white-matter tissue and hippocampus. However, while the method of Dey *et al*. performed better than other methods at tracking the distribution of cortical grey-matter volume, there was still noticeable underestimation. The methods of Dalca *et al*. and Dey *et al*. were able to track the hippocampal volumes with good accuracy, and although they were the best for cortical grey-matter out of the methods compared here, they were still only capturing about half of the empirically observed change in volume. For the ventricles and white-matter, the method of Dalca *et al*. performed poorly.

Another output of interest from these methods are the template images themselves and how suitable they are for registration targets and how well they capture a qualitative representation of the ageing process. For the method of Yu *et al*., conditional probabilistic templates were produced that have a number of practical uses, including the tracking of various morphological features, such as structural volumes; however, further work is required to enable such an approach to effectively produce T1w structural templates that appear realistic and would make good targets for registrations. In contrast, the method of Dalca *et al*. employed a relatively unconstrained approach in their first network stage, with the network able to make independent voxel-wise intensity changes to the conditional template images. As such, this kind of architecture can introduce intensity artefacts into the conditional templates and lacks the strong underpinning for finding anatomical correspondences across the age range of a method based on spatial deformations. Despite this, the method of Dey *et al*., while employing a similarly unconstrained approach in their first network stage, was able to produce conditional templates with fewer intensity artefacts, although this method still exhibits negative Jacobian determinants in deformations between conditional templates and underlying participant scans, and similarly lacks the same underpinning as a method based purely on spatial deformations for finding anatomical correspondences across the age range. The use of a diffeomorphic deformation framework in both stage 1 and 2 of our network means that conditional templates will typically have few artefacts introduced, with it focusing on geometric changes of interest.

This work demonstrates that a key advantage of a purely geometric method, that uses a diffeomorphic deformation framework throughout the network architecture, is that it is able to provide a geometric link, in a topologically consistent manner, between common reference templates or standard atlases, conditional templates, and participant scans. Such geometric linkage is important in many neuroimaging studies, as it allows for more nuanced comparisons that account for various demographic and clinical variables of interest while empowering effective comparisons across studies that utilise a common reference frame or standard atlases. Moreover, this may facilitate cross-study comparisons that foster improvements in the calibration of treatments and interventions in personalised medicine.

### 4.1 Limitations

One limitation of our method is the inability of our network to make non-geometric changes to conditional templates as conditioning parameters, such as age, are adjusted. For instance, it is known that during ageing the human brain goes through a process of demyelination [47, 48], leading to an increase in T1-relaxation time for the affected tissue, and hence decreased signal intensity in T1-weighted images. These kinds of changes cannot easily be captured by a purely geometric network architecture. We have investigated the inclusion of network modules that are able to make non-geometric changes to conditional templates and predicted participant scans, however, we found that they made the model unstable and less able to capture geometric changes of interest. One approach to address this issue that may be worth investigating is to model non-geometric, voxel-wise changes of intensities such that these changes are constrained to stay independent of the geometric changes and do not impact geometric adjustments associated with the shift in anatomical borders.

Another limitation of our approach, and its potential, lies in the ability of local mean constraints to be effectively scaled-up to use with higher-dimensional combinations of conditional parameters. Given that our method partitioned the conditional parameter space to create a discrete number of voxel-wise mean-tracking deformations, the combinatorial increase in the number of deformations required with higher dimensions may become difficult to manage computationally. For instance, if we were to jointly model 4 conditioning parameters, each partitioned such that we were tracking 10 discrete representations for each conditioning parameter of interest, we would need to manage 10^4^ local mean deformations if they were all allowed to be independent. Additionally, the amount of data used to inform each mean deformation becomes smaller in higher dimensions due to data sparsity. The first issue may be ameliorated by managing relevant data-structures on disk or in CPU memory, as opposed to GPU memory. However, the sparsity of participant scans in the conditional space means that either different representations over the conditional parameter range are needed, which might be coarser or better at modeling/interpolating, or that additional training data is required to reduce the sparsity. Relaxing the independence assumption and imposing a better model is something that deep learning may be able to provide, although the exact form of the model and the amount of bias that it induces would require careful management. While the methods of Dalca *et al*. and Dey *et al*. utilised a global mean-tracking constraint across the dataset, which has some benefits to encourage centrality, such a global constraint may suffer from bias towards particular parts of conditional parameter space. For instance, this kind of network constraint is likely to become highly optimised for the most common conditioning parameters.

While we have prepared our structural volumetric analysis for four structures that are strongly implicated in ageing and neurodegeneration, and also commonly reported on, there are many others that may be useful. Hence the structures that are considered can be diversified and expanded in future studies. Additionally, the calculation of volumes used here was based on the output from an automated segmentation tool, albeit one with proven performance, but it is not known whether these segmentations are biased or creating errors that may impact these results. This may be addressed in future studies by employing additional segmentation methods to establish the degree to which they produce consistent volumetric outputs for a given conditional template.

Another consideration is the size of the dataset utilised for this study, namely 1534 scans across 515 individual participants. It is possible that some or all of the methods that we have evaluated perform significantly better on much larger datasets. However, the use of a more constrained dataset has certain advantages for our evaluation: (i) some populations, especially of rarer diseases that might be of interest, will necessitate working with smaller datasets; and when introducing multiple conditioning factors, the density of datapoints in the higher dimensional space will decrease — hence, methods will need to be able to cope in regimes where the density of datapoints is lower than the density of datapoints found in large single-factor datasets. Despite this, expanding the size of the dataset, and in particular, the number of independent participants, in future work, may assist in establishing the effectiveness of these methods in settings where datapoints are less constrained.

### 4.2 Future directions and conclusion

An interesting direction for future work could be to mix the different elements from the methods compared here. For instance, the method of Yu *et al*. used probabilistic atlases, which could potentially be leveraged in conjunction with intensity templates to provide additional information to a network when producing T1-weighted conditional templates. Here, we can envisage probabilistic atlases augmenting templates to provide richer information about key brain structures in the network. Additionally, the use of adversarial training in the method of Dey *et al*. has enabled this method to produce conditional templates that are better calibrated across a range of criteria when compared to the method of Dalca *et al*. — as such, this training approach may prove beneficial in future works. Furthermore, while we did not see improvements to conditional template generation for our method when using an NCC reconstruction loss and FiLM conditioning, it may prove beneficial to use these modelling techniques in future works that blend aspects of the various methods considered in this work.

In conclusion, while our approach, and those of Dalca *et al*., Yu *et al*., and Dey *et al*. offer a number of useful contributions to conditional template construction, current deep-learning methods require further refinement to achieve stronger calibration with morphological changes exhibited by ground truth data in ageing and degenerative brains. Furthermore, when considering building methods that can work with a number of conditional parameters jointly, conventional methods often have limitations in terms of the large amount of data required or the restricted modeling options. Hence, deep-learning approaches have some inherent advantages in terms of flexible datadriven modeling, without necessarily requiring as much data, and so future work in this area should be pursued while actively working to improve the ability of these methods to accurately track volumetric changes while maintaining other desirable characteristics, such as topological consistency.

## Acknowledgements

Data collection and sharing for this project was funded by the Alzheimer’s Disease Neuroimaging Initiative (ADNI) (National Institutes of Health Grant U01 AG024904) and DOD ADNI (Department of Defense award number W81XWH-12-2-0012). ADNI is funded by the National Institute on Aging, the National Institute of Biomedical Imaging and Bioengineering, and through generous contributions from the following: AbbVie, Alzheimer’s Association; Alzheimer’s Drug Discovery Foundation; Araclon Biotech; BioClinica, Inc.; Biogen; Bristol-Myers Squibb Company; CereSpir, Inc.; Cogstate; Eisai Inc.; Elan Pharmaceuticals, Inc.; Eli Lilly and Company; EuroImmun; F. Hoffmann-La Roche Ltd and its affiliated company Genentech, Inc.; Fujirebio; GE Healthcare; IXICO Ltd.; Janssen Alzheimer Immunotherapy Research & Development, LLC.; Johnson & Johnson Pharmaceutical Research & Development LLC.; Lumosity; Lundbeck; Merck & Co., Inc.; Meso Scale Diagnostics, LLC.; NeuroRx Research; Neurotrack Technologies; Novartis Pharmaceuticals Corporation; Pfizer Inc.; Piramal Imaging; Servier; Takeda Pharmaceutical Company; and Transition Therapeutics. The Canadian Institutes of Health Research is providing funds to support ADNI clinical sites in Canada. Private sector contributions are facilitated by the Foundation for the National Institutes of Health (www.fnih.org). The grantee organization is the Northern California Institute for Research and Education, and the study is coordinated by the Alzheimer’s Therapeutic Research Institute at the University of Southern California. ADNI data are disseminated by the Laboratory for Neuro Imaging at the University of Southern California.

## Footnotes

1 voxelmorph.csail.mit.edu (downloaded 23 Jul 2023)

2 github.com/evanmy/conditional_deformation (downloaded 1 Nov 2023)

3 github.com/neel-dey/Atlas-GAN (downloaded 4 Sep 2024)

