## Supplementary materials for "Deep-diffeomorphic networks for conditional brain templates"

### Deep-diffeomorphic networks for conditional brain templates: supplementary materials

#### S1 Structural Similarity Index Measure (SSIM) derivation

SSIM extracts and calculates comparisons for three key images features, namely: (i) luminance, $l(\cdot)$; (ii) contrast, $c(\cdot)$; and (iii) structure, $s(\cdot)$; combining the components to produce an overall similarity measure for images $\mathbf{x}$ and $\mathbf{y}$: $S(\mathbf{x},\mathbf{y}) = f(l(\mathbf{x},\mathbf{y}), c(\mathbf{x},\mathbf{y}), s(\mathbf{x},\mathbf{y}))$.

The luminance comparison function, $l(\mathbf{x},\mathbf{y})$, for images $\mathbf{x}$ and $\mathbf{y}$, compares the mean pixel or voxel intensities, $\mu_{\mathbf{x}}$ and $\mu_{\mathbf{y}}$, for two images, as follows:

$$l(\mathbf{x},\mathbf{y}) = \frac{2\mu_{\mathbf{x}}\mu_{\mathbf{y}} + C1}{\mu_{\mathbf{x}}^2 + \mu_{\mathbf{y}}^2 + C1} \tag{1}$$

where $C1 = (K_1 L)^2$ is a constant term for numerical stability, $L$ is the pixel or voxel dynamic range and $K_1 << 1$ is a small constant.

The contrast comparison function, $c(\mathbf{x},\mathbf{y})$, takes a similar form to $l(\cdot)$, comparing the standard deviation of the intensity values taken across all voxels within an image, $\sigma_{\mathbf{x}}$ and $\sigma_{\mathbf{y}}$, for two images, as follows:

$$c(\mathbf{x},\mathbf{y}) = \frac{2\sigma_{\mathbf{x}}\sigma_{\mathbf{y}} + C2}{\sigma_{\mathbf{x}}^2 + \sigma_{\mathbf{y}}^2 + C2} \tag{2}$$

where $C2$ is similarly calculated to $C1$ for $K_2 << 1$.

To perform a structural comparison between images $\mathbf{x}$ and $\mathbf{y}$, luminance is subtracted from the images and they are normalised by variance (i.e., $(\mathbf{x} - \mu_{\mathbf{x}})/\sigma_{\mathbf{x}}$ and $(\mathbf{y} - \mu_{\mathbf{y}})/\sigma_{\mathbf{y}}$), with the correlation between these an effective measure to quantify structural similarity, as given by:

$$s(\mathbf{x},\mathbf{y}) = \frac{\sigma_{\mathbf{xy}} + C3}{\sigma_{\mathbf{x}}\sigma_{\mathbf{y}} + C3} \tag{3}$$

where $\sigma_{\mathbf{xy}}$ represents the covariance of the intensity values taken across all voxels in images $\mathbf{x}$ and $\mathbf{y}$. Each of the three components is then weighted to give an overall measure of SSIM:

$$S(\mathbf{x},\mathbf{y}) = [l(\mathbf{x},\mathbf{y})]^{\alpha} \cdot [c(\mathbf{x},\mathbf{y})]^{\beta} \cdot [s(\mathbf{x},\mathbf{y})]^{\gamma} \tag{4}$$

where $\alpha, \beta, \gamma > 0$ adjust the importance of each component. Typically, each of these exponents is set equal to 1; which we have adopted for the purposes of our analysis.

#### S2 Distributions of Jacobian determinants for models estimated using reduced data sample-sizes or corrupted data

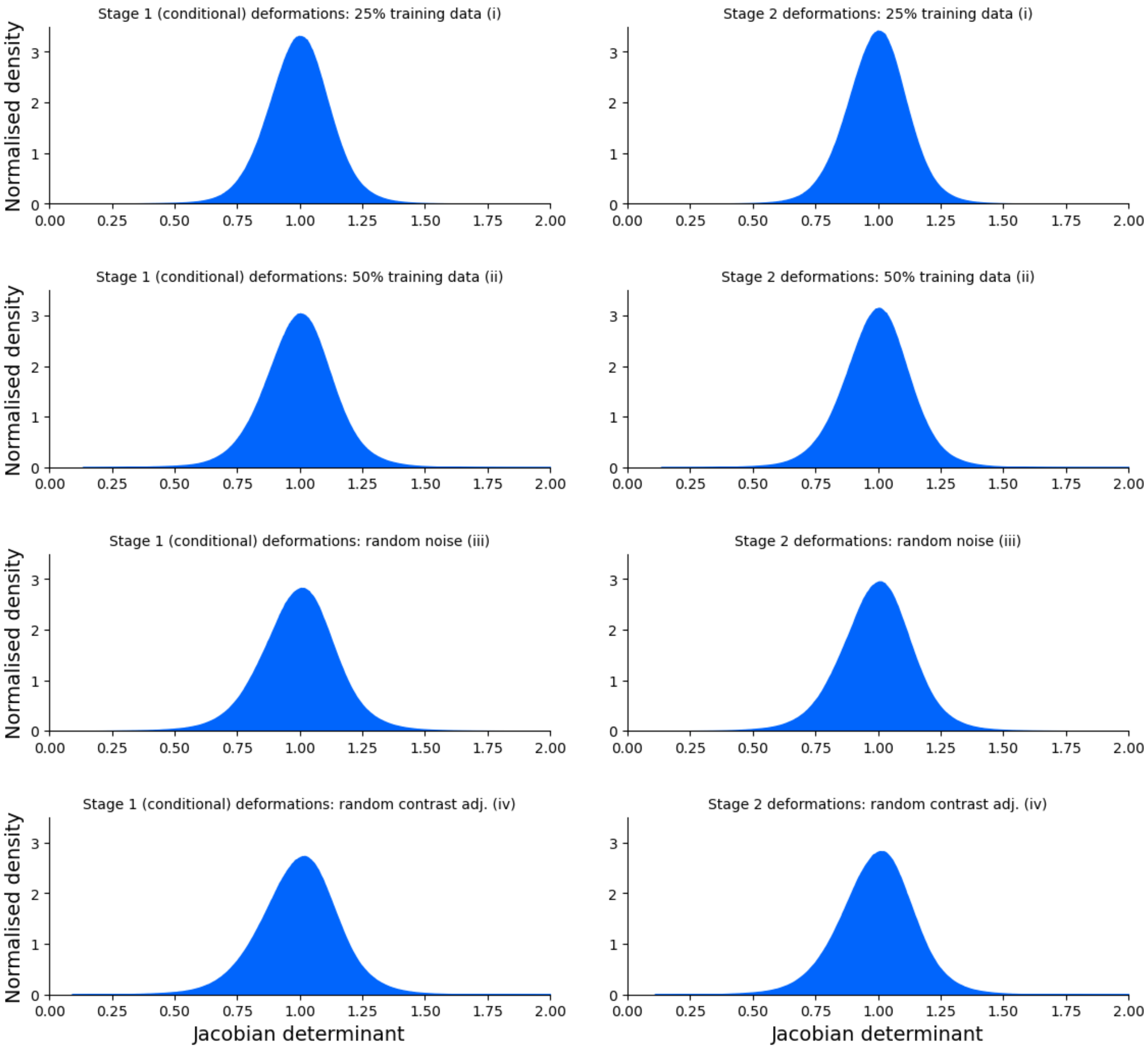


**Figure 1:** *Distributions of Jacobian determinants for our method trained on perturbed training datasets for deformation fields from unconditional to conditional templates, and conditional templates to predicted participant scans: (i) using 25% of the training dataset, randomly sampled; (ii) using 50% of the training dataset, randomly sampled; (iii) adding random Gaussian noise to normalised training dataset scans (i.e., normalised by the value of the 99.9$^{th}$ percentile and clipped to a maximum intensity value of 1.1) with a standard deviation of 0.03; and (iv) randomly adjusting the contrast of training scans by raising each scan's voxels to $\gamma$, where $\gamma = e^{\beta}$, $\beta \sim \mathcal{U}(-a, a)$, and a is set to 0.3.*

#### S3 Structural similarity: SSIM and RMSD for models estimated using reduced data sample-sizes or corrupted data

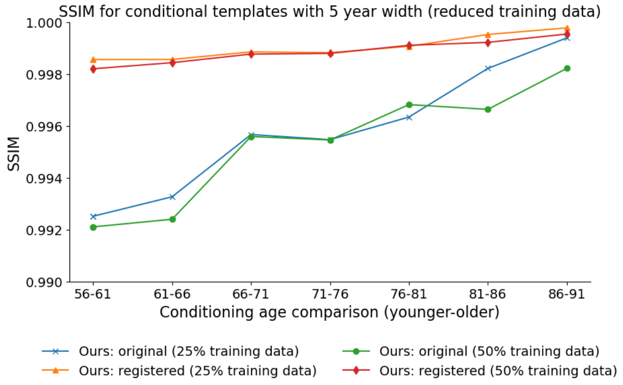


**(a)** *SSIM: reduced training data*

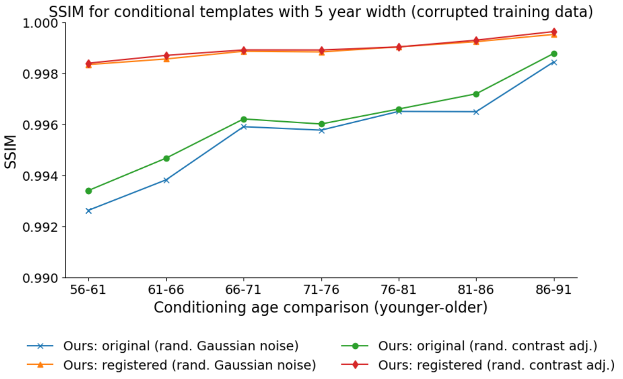


**(b)** *SSIM: corrupted training data*

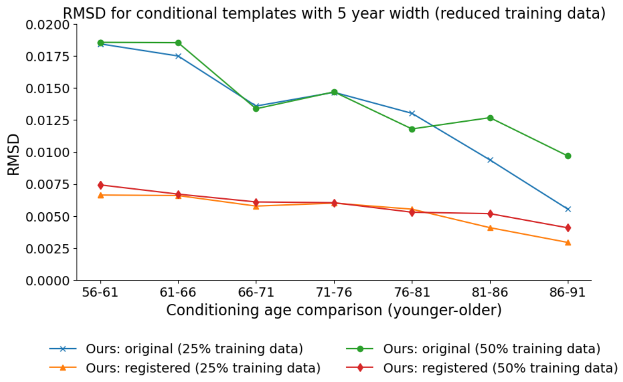


**(c)** *RMSD: reduced training data*

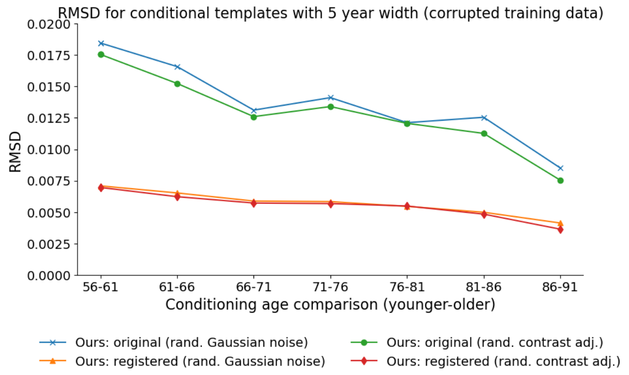


**(d)** *RMSD: corrupted training data*

**Figure 2:** ***(Top)*** *Structural Similarity Index Measure (SSIM), and* ***(Bottom)*** *root mean squared difference (RMSD) between estimated conditional templates separated by 5 years, on both a pre- (original) and post-registration basis. Our method is trained on:* ***(a, c)*** *25% and 50% of the full training dataset (as described in section 2.3), and* ***(b, d)*** *training data that is corrupted with random Gaussian noise or random contrast adjustments. For the registered images, each lower-age conditional template has been registered to the next higher age conditional template, with the registered image and the higher age conditional template used to calculate SSIM and RMSD on a post-registration basis.*

#### S4 Volumetric measurements for models estimated using reduced data sample-sizes or corrupted data

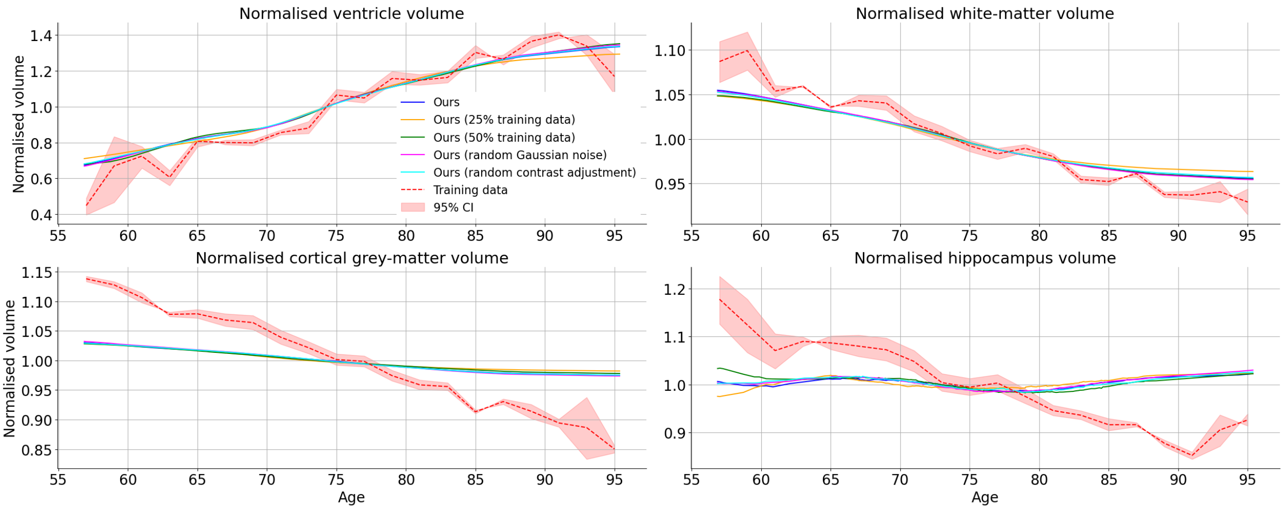


**Figure 3:** *Volumetric measurements for various ROIs where the training set has been perturbed: cortical grey-matter, white-matter, lateral ventricles and hippocampus. We have perturbed the training set in various ways: (i) randomly sampling 25% of the training dataset; (ii) randomly sampling 50% of the training dataset; (iii) adding random Gaussian noise to normalised training dataset scans (i.e., normalised by the value of the 99.9$^{th}$ percentile and clipped to a maximum intensity value of 1.1) with a standard deviation of 0.03; and (iv) randomly adjusting the contrast of training scans by raising each scan's voxels to $\gamma$, where $\gamma = e^{\beta}$, $\beta \sim \mathcal{U}(-a, a)$, and a is set to 0.3. Volumes are presented on a normalised basis, where volumes are normalised by the mean of all volumes across the range of conditional parameters, independently for each ROI and method. In addition, a 95% confidence interval is also presented for the empirical estimates made from training data within 2-year age bins.*

#### S5 Distributions of Jacobian determinants for models estimated using two covariates (age and sex)

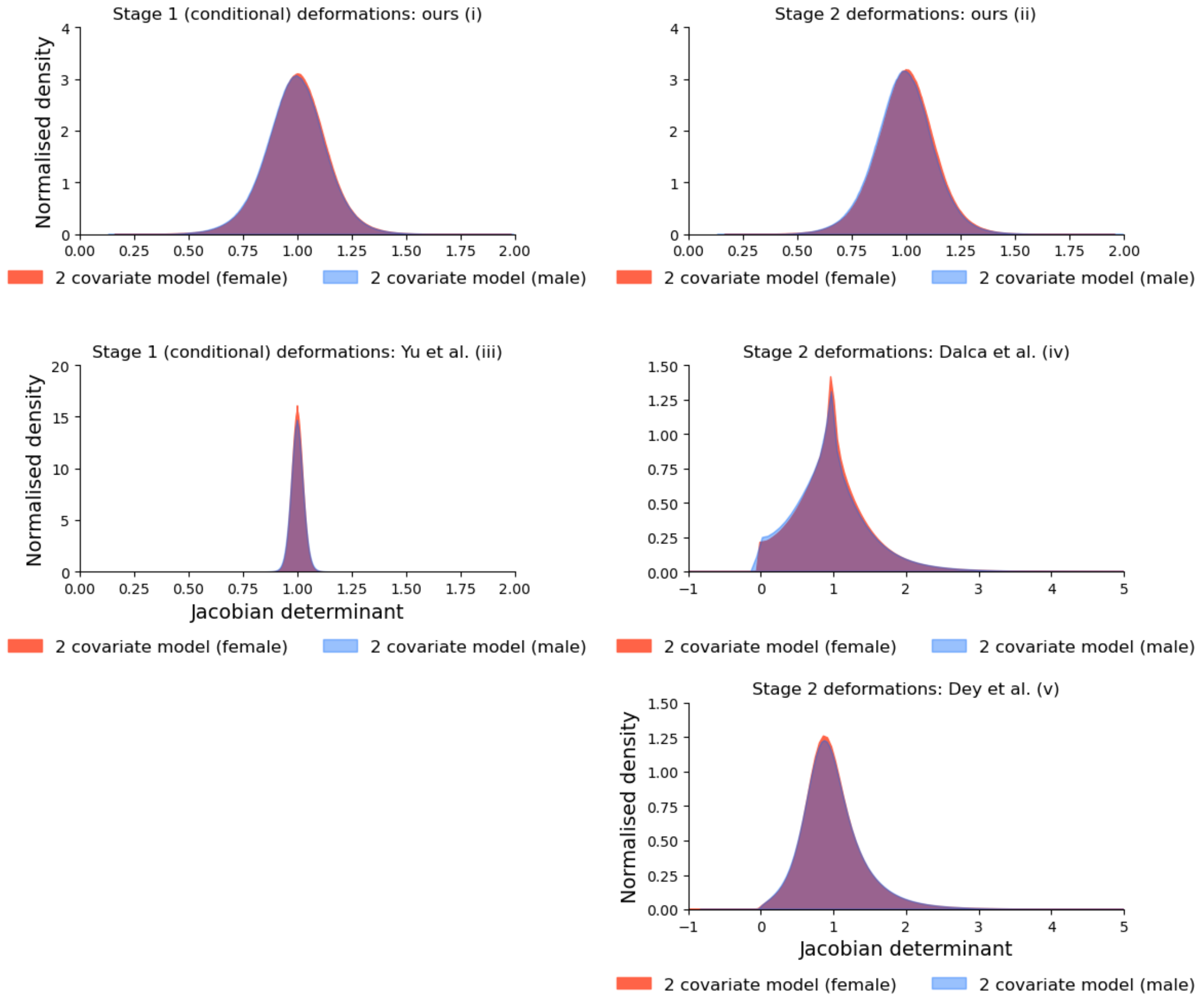


**Figure 4:** *Distributions of Jacobian determinants for models trained on the full training dataset with two covariates (age and sex) used to generate conditional templates, for: (i) our method for deformation fields from unconditional to conditional templates, and (ii) conditional templates to predicted participant scans; (iii) deformations from unconditional to conditional templates for the method of Yu et al.; (iv) deformations from conditional templates to predicted participant scans for the method of Dalca et al.; (v) and deformations from conditional templates to predicted participant scans for the method of Dey et. al.*

#### S6 Structural similarity: SSIM and RMSD for models estimated using two covariates (age and sex)

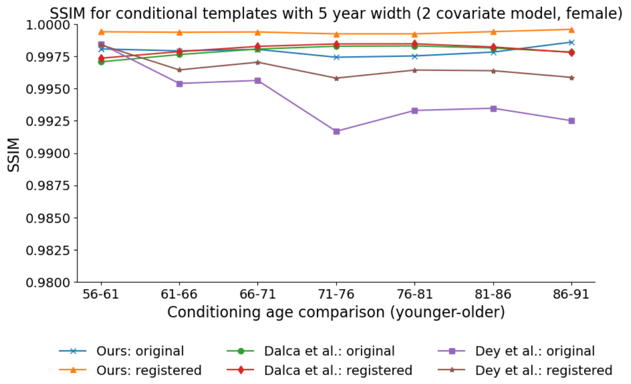


**(a)** *SSIM: 2 covariate model, female*

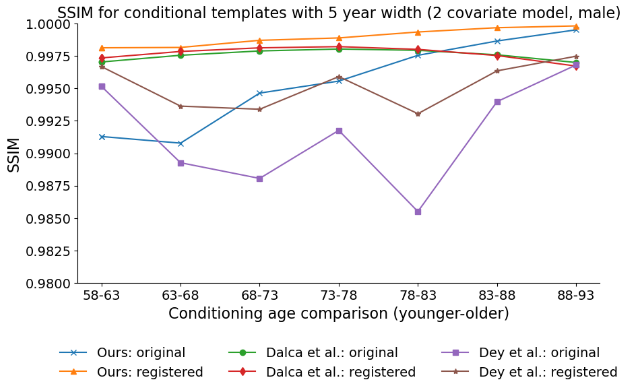


**(b)** *SSIM: 2 covariate model, male*

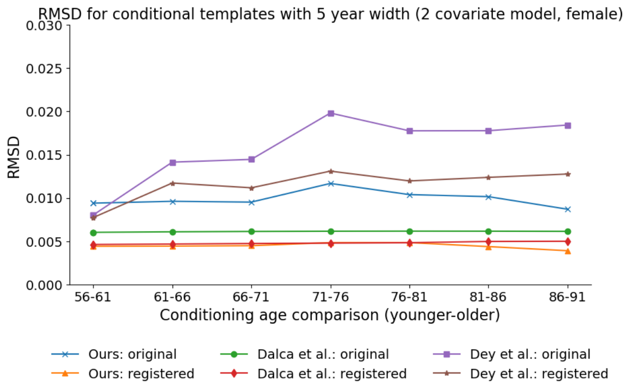


**(c)** *RMSD: 2 covariate model, female*

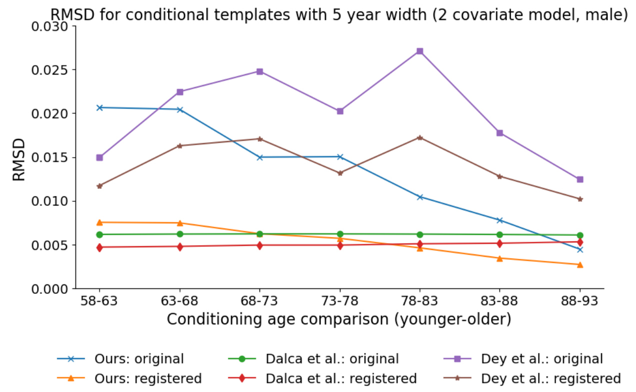


**(d)** *RMSD: 2 covariate model, male*

**Figure 5:** ***(Top)*** *Structural Similarity Index Measure (SSIM), and* ***(Bottom)*** *root mean squared difference (RMSD) between estimated conditional templates separated by 5 years, on both a pre- (original) and post-registration basis. Our method and the methods of Dalca et. al. and Dey et. al. are trained on the full training dataset with two covariates (age and sex) used to generate conditional templates. For the registered images, each lower-age conditional template has been registered to the next higher age conditional template, with the registered image and the higher age conditional template used to calculate SSIM and RMSD on a post-registration basis.*

### S7 Volumetric measurements for models estimated using two covariates (age and sex)

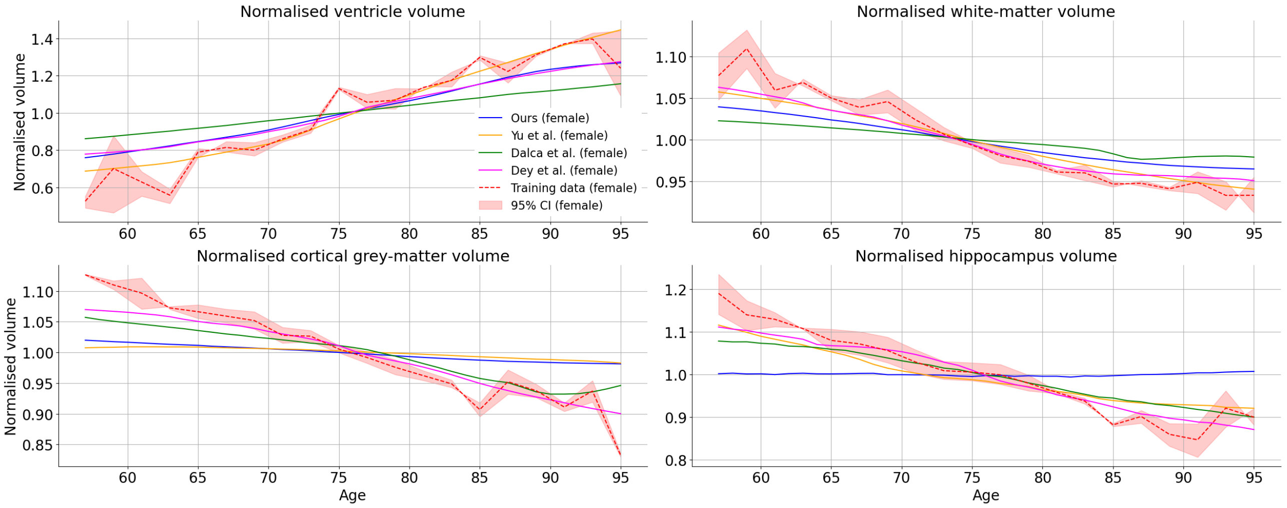


**(a)** *Normalised volume: 2 covariate model, female*

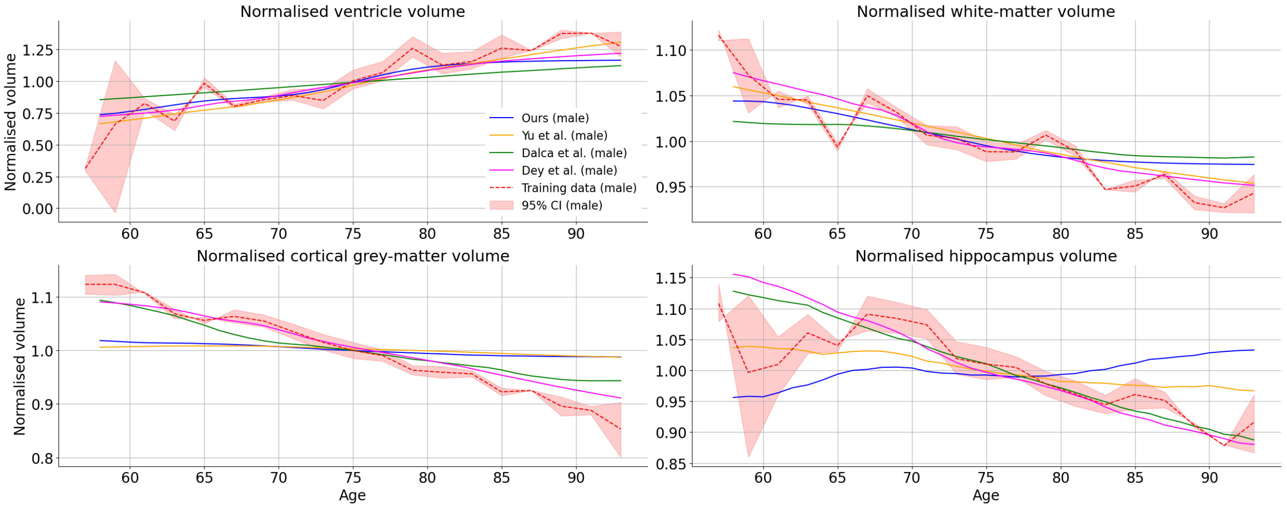


**(b)** *Normalised volume: 2 covariate model, male*

**Figure 6:** *Volumetric measurements for various ROIs: cortical grey-matter, white-matter, lateral ventricles and hippocampus. Each method was trained on the full training dataset with two covariates (age and sex) used to generate conditional templates. Volumes are presented on a normalised basis, where volumes are normalised by the mean of all volumes across the range of conditional parameters, independently for each ROI and method. In addition, a 95% confidence interval is also presented for the empirical estimates made from training data (female) within 2-year age bins.*

#### S8 Distributions of Jacobian determinants for models estimated using female- and male-only data

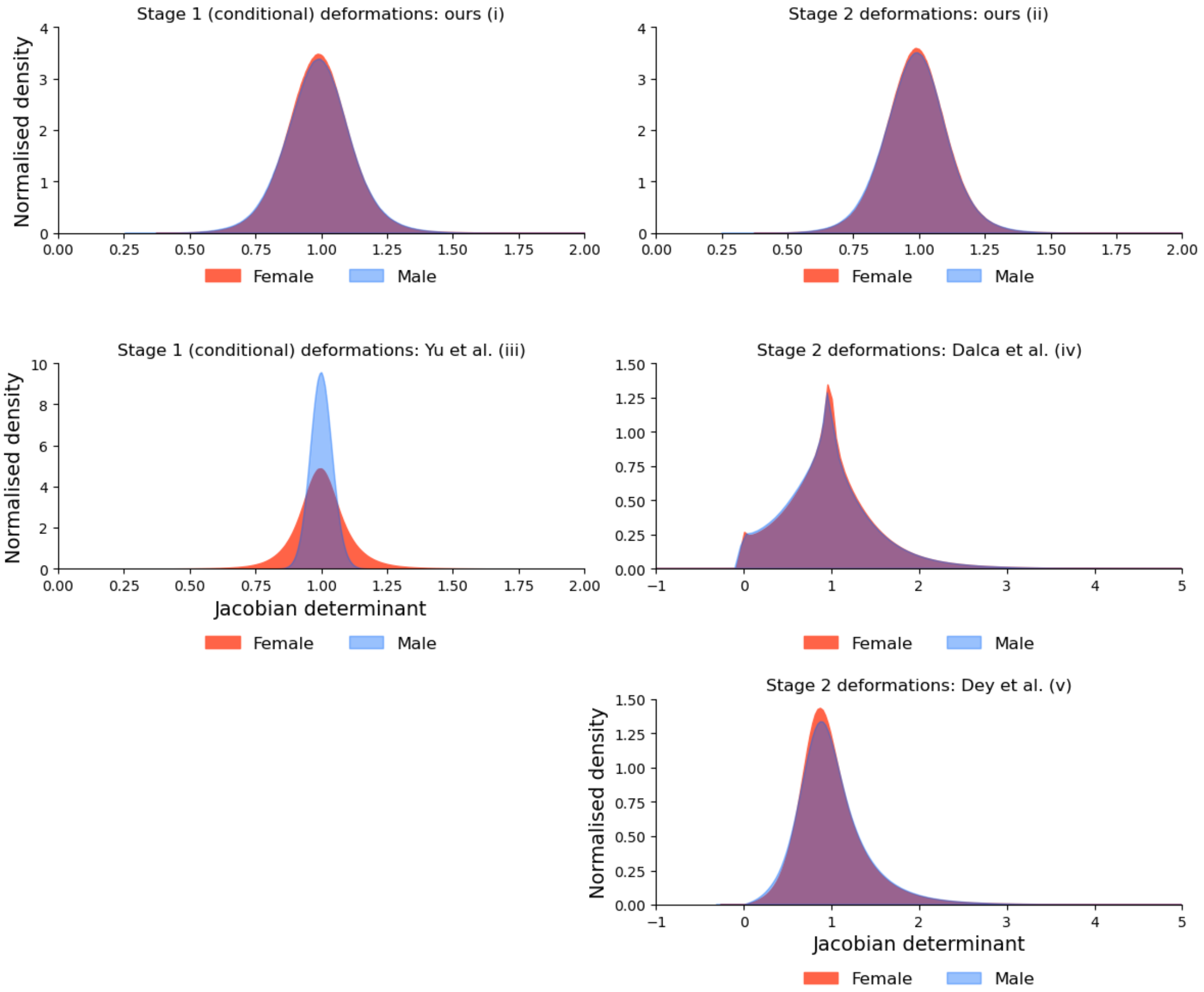


**Figure 7:** *Distributions of Jacobian determinants for models trained on a female- and male-only dataset for: (i) our method for deformation fields from unconditional to conditional templates, and (ii) conditional templates to predicted participant scans; (iii) deformations from unconditional to conditional templates for the method of Yu et al.; (iv) deformations from conditional templates to predicted participant scans for the method of Dalca et al.; (v) and deformations from conditional templates to predicted participant scans for the method of Dey et. al.*

#### S9 Structural similarity: SSIM and RMSD for models estimated on female- and male-only data

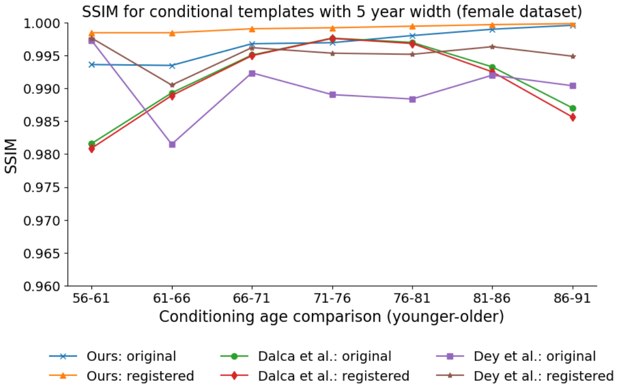


**(a)** *SSIM: female data*

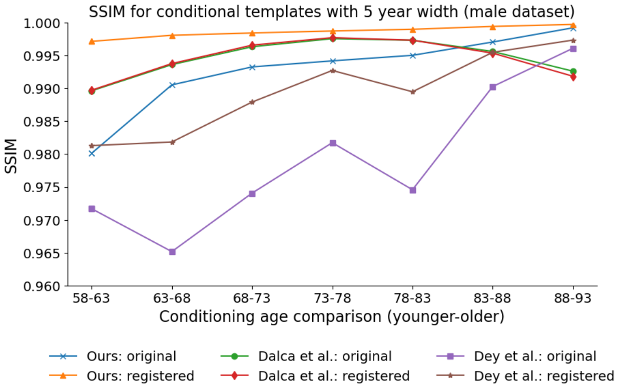


**(b)** *SSIM: male data*

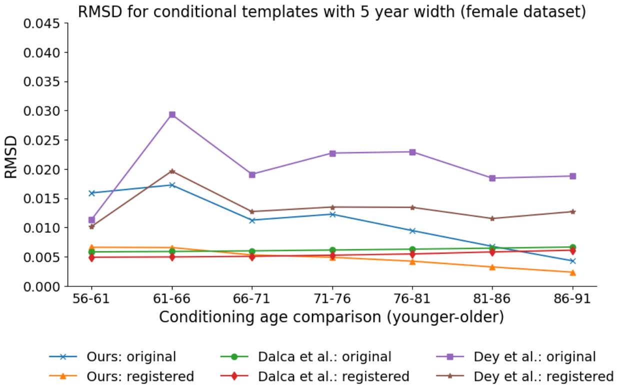


**(c)** *RMSD: female data*

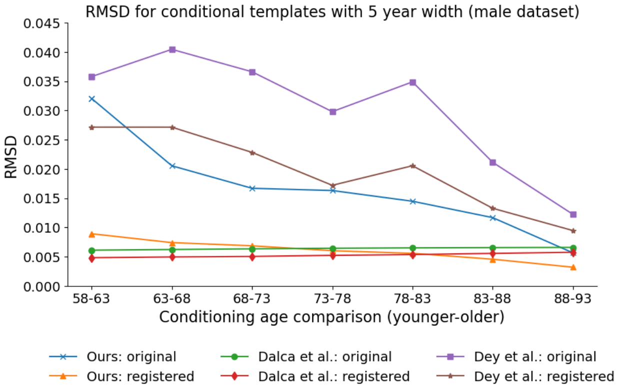


**(d)** *RMSD: male data*

**Figure 8:** ***(Top)*** *Structural Similarity Index Measure (SSIM), and* ***(Bottom)*** *root mean squared difference (RMSD) between estimated conditional templates separated by 5 years, on both a pre- (original) and post-registration basis. Our method and the methods of Dalca et. al. and Dey et. al. are trained on:* ***(a, c)*** *a female-only partition of the training dataset (as described in section 2.3), and* ***(b, d)*** *the corresponding male-only partition. For the registered images, each lower-age conditional template has been registered to the next higher age conditional template, with the registered image and the higher age conditional template used to calculate SSIM and RMSD on a post-registration basis.*

#### S10 Volumetric measurements for models estimated using female- and male-only data

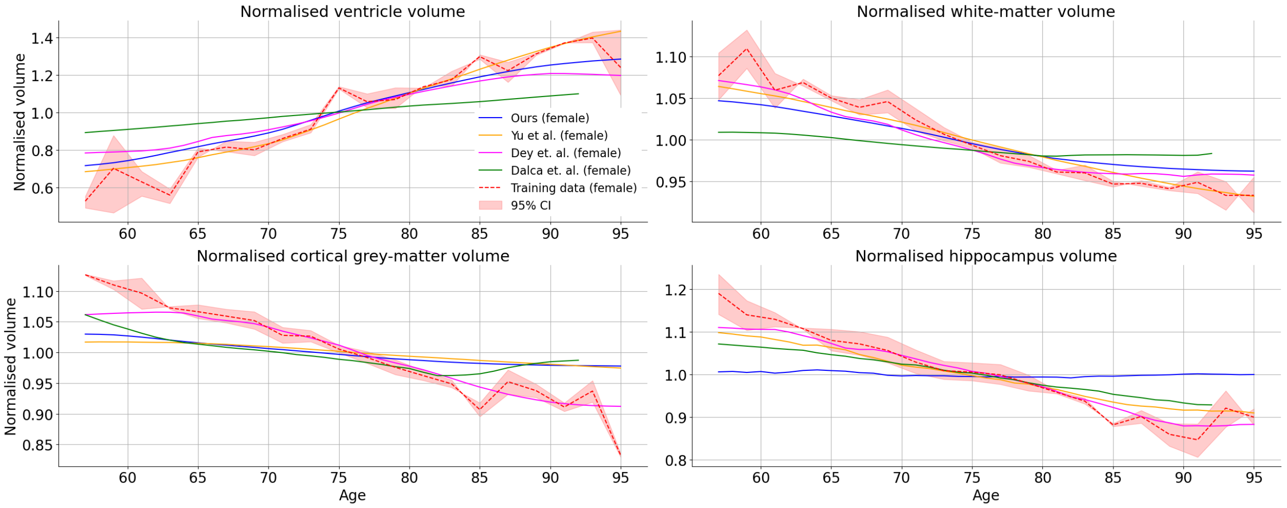


**(a)** *Normalised volume: female data*

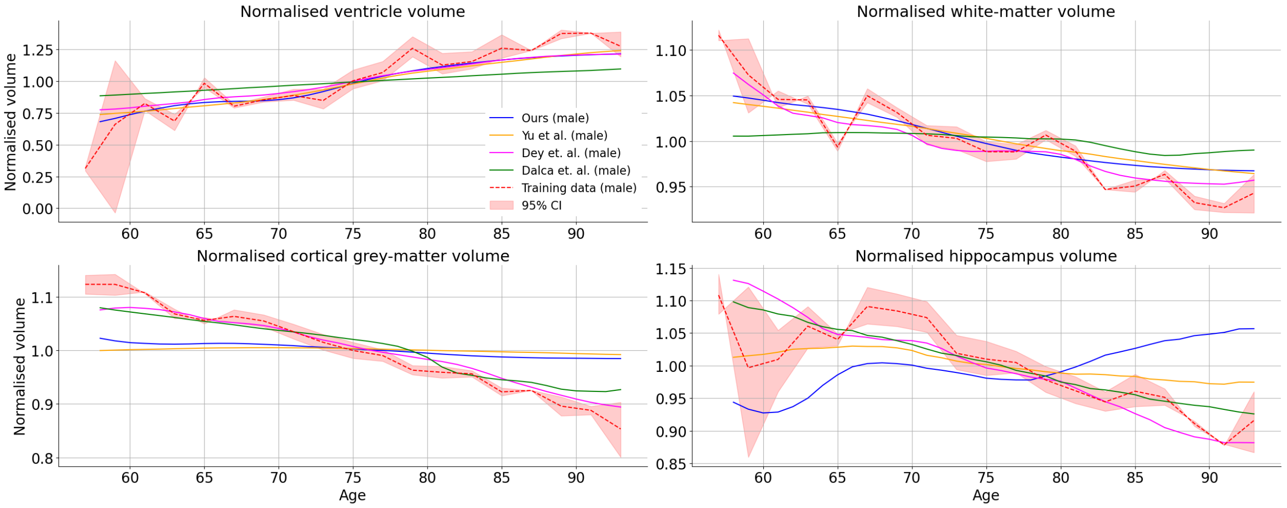


**(b)** *Normalised volume: male data*

**Figure 9:** *Volumetric measurements for various ROIs: cortical grey-matter, white-matter, lateral ventricles and hippocampus. Each method was trained on:* ***(a)*** *a female-only partition of the training dataset (as described in section 2.1), and* ***(b)*** *the corresponding male-only partition. Volumes are presented on a normalised basis, where volumes are normalised by the mean of all volumes across the range of conditional parameters, independently for each ROI and method. In addition, a 95% confidence interval is also presented for the empirical estimates made from training data (female) within 2-year age bins.*

#### S11 Distributions of Jacobian determinants for models estimated with loss function components iteratively set to zero

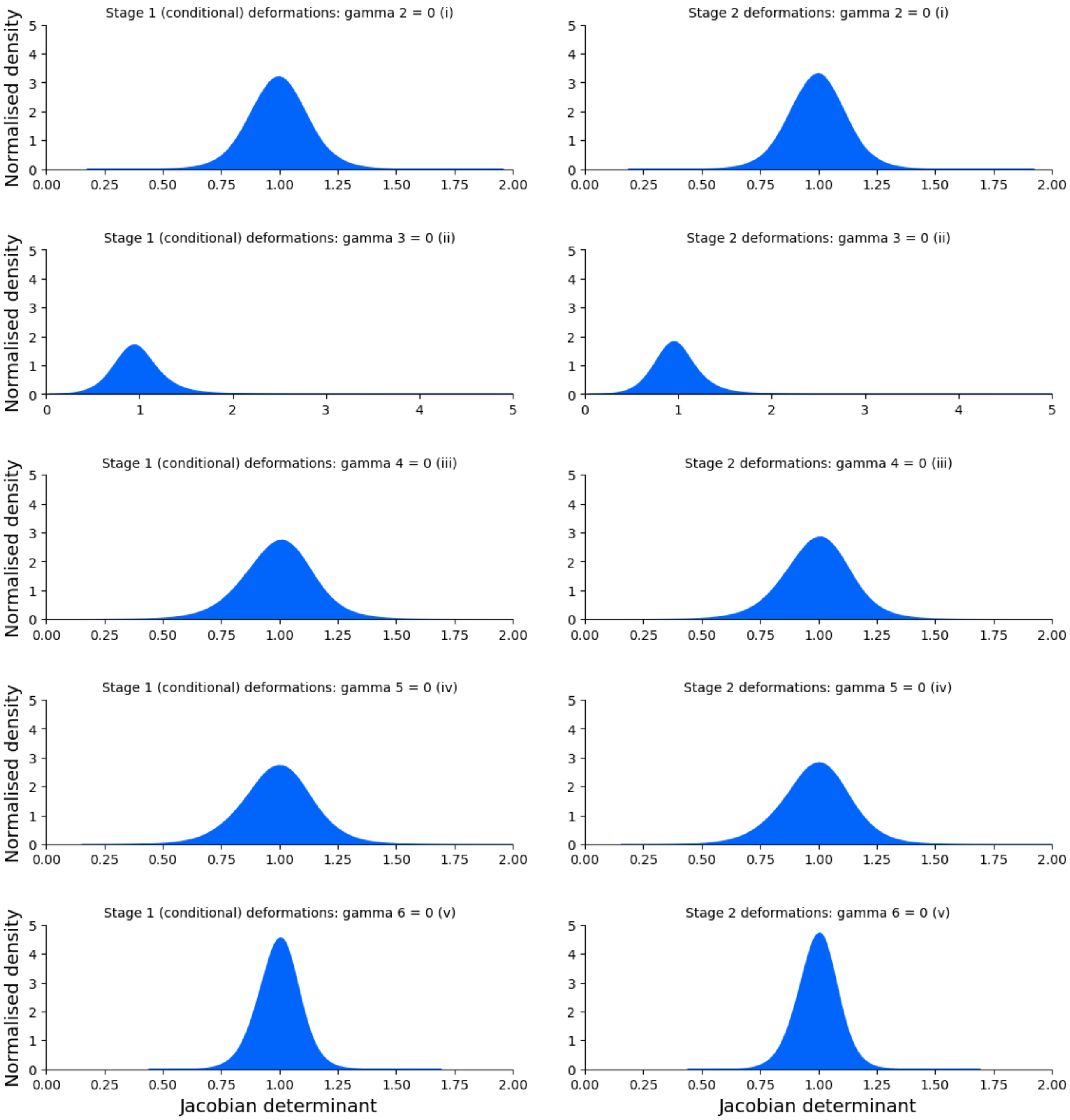


**Figure 10:** *Distributions of Jacobian determinants for our method when loss function components, $\gamma_2$–$\gamma_6$ are iteratively set to zero: (i) $\gamma_2 = 0$, (ii) $\gamma_3 = 0$, (iii) $\gamma_4 = 0$, (iv) $\gamma_5 = 0$, and (v) $\gamma_6 = 0$.*

#### S12 Structural similarity: SSIM and RMSD for models estimated with loss function components iteratively set to zero

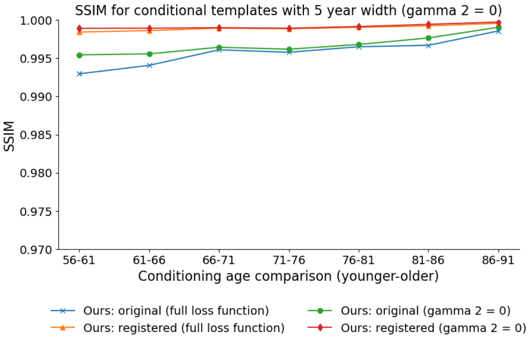


**(a)** *SSIM:* $\gamma_2 = 0$

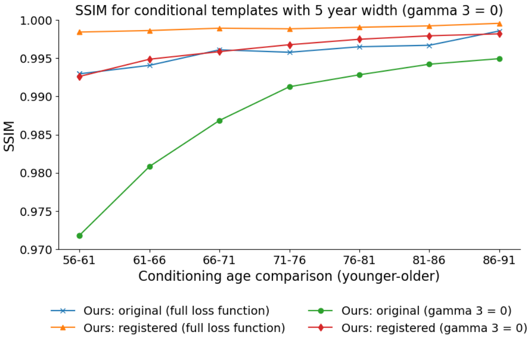


**(b)** *SSIM:* $\gamma_3 = 0$

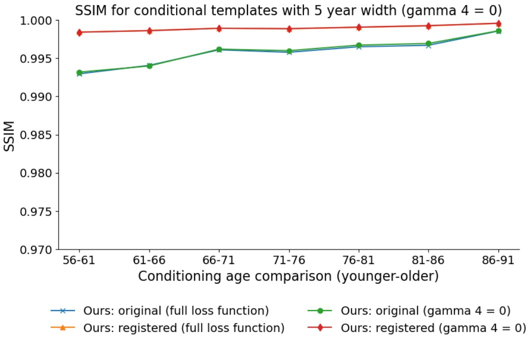


**(c)** *SSIM:* $\gamma_4 = 0$

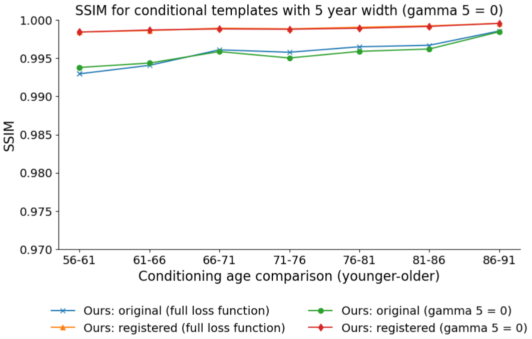


**(d)** *SSIM:* $\gamma_5 = 0$

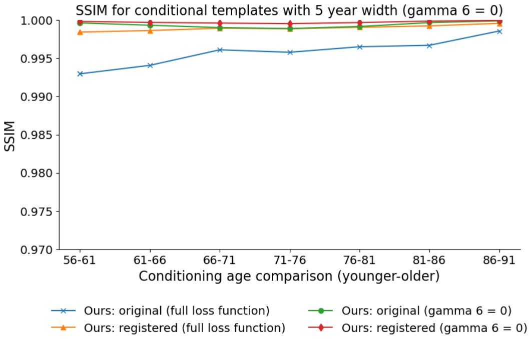


**(e)** *SSIM:* $\gamma_6 = 0$

**Figure 11:** *Structural Similarity Index Measure (SSIM) between estimated conditional templates separated by 5 years, on both a pre- (original) and post-registration basis, where:* $\gamma_2$*–*$\gamma_6$ *are iteratively set to zero. For comparative purposes, we have provided SSIM calculations for our method where the full loss function is used. For the registered images, each lower-age conditional template has been registered to the next higher age conditional template, with the registered image and the higher age conditional template used to calculate SSIM on a post-registration basis.*

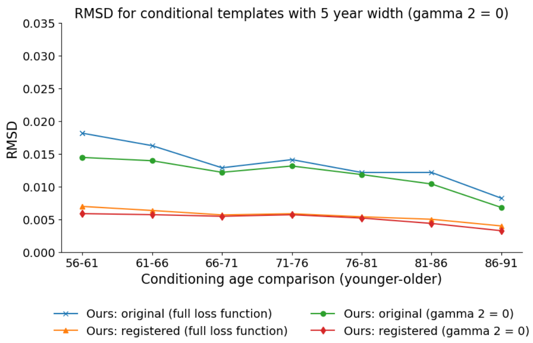


**(a)** *RMSD:* $\gamma_2 = 0$

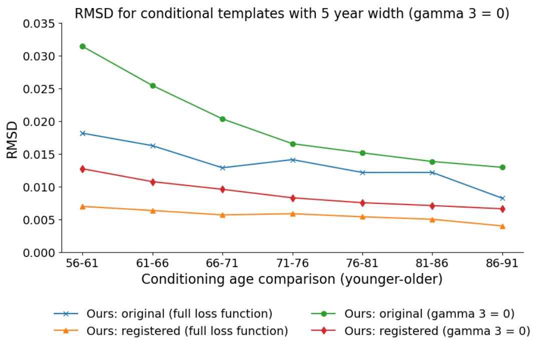


**(b)** *RMSD:* $\gamma_3 = 0$

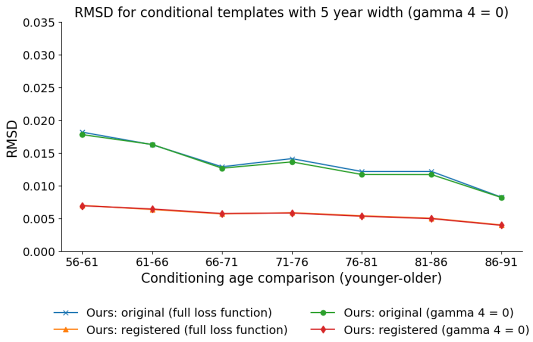


**(c)** *RMSD:* $\gamma_4 = 0$

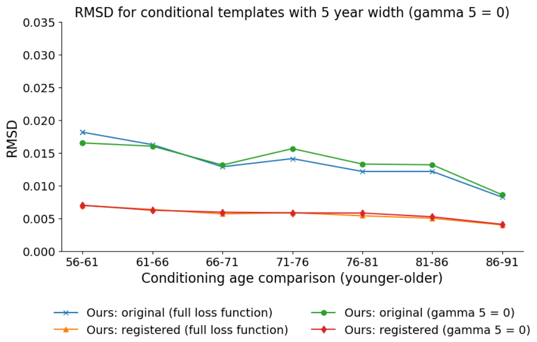


**(d)** *RMSD:* $\gamma_5 = 0$

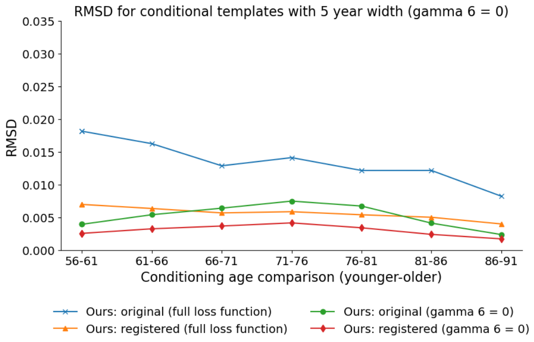


**(e)** *RMSD:* $\gamma_6 = 0$

**Figure 12:** *Root mean squared difference (RMSD) between estimated conditional templates separated by 5 years, on both a pre- (original) and post-registration basis, where:* $\gamma_2$–$\gamma_6$ *are iteratively set to zero. For comparative purposes, we have provided RMSD calculations for our method where the full loss function is used. For the registered images, each lower-age conditional template has been registered to the next higher age conditional template, with the registered image and the higher age conditional template used to calculate RMSD on a post-registration basis.*

#### S13 Volumetric measurements for models estimated with loss function components iteratively set to zero

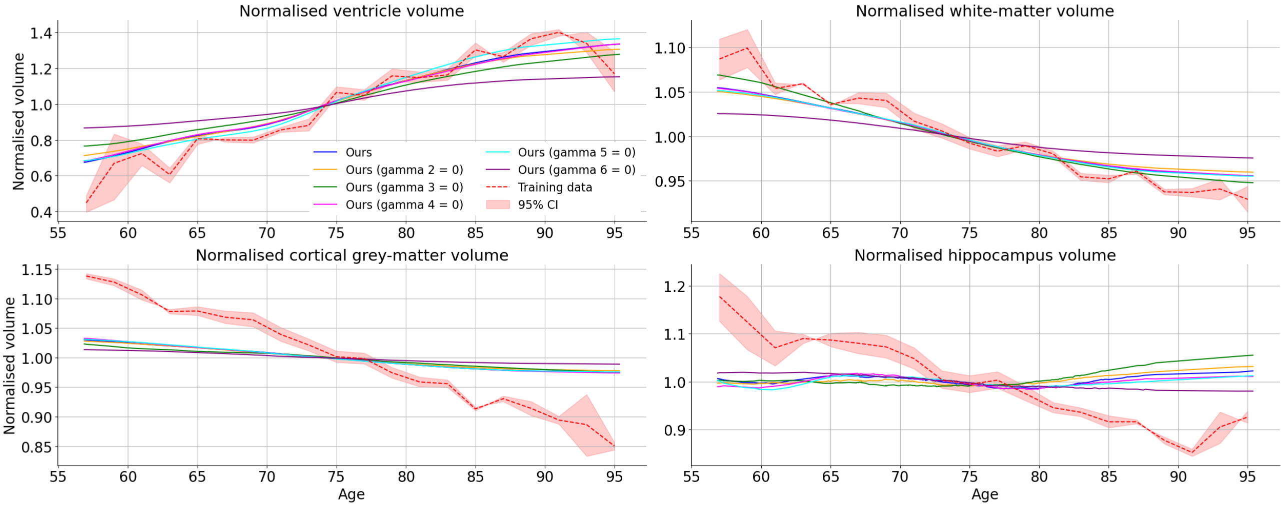


**Figure 13:** *Volumetric measurements for various ROIs where the training set has been perturbed: cortical grey-matter, white-matter, lateral ventricles and hippocampus. We have perturbed the training set in various ways: (i) randomly sampling 25% of the training dataset; (ii) randomly sampling 50% of the training dataset; (iii) adding random Gaussian noise to normalised training dataset scans (i.e., normalised by the value of the 99.9$^{th}$ percentile and clipped to a maximum intensity value of 1.1) with a standard deviation of 0.03; and (iv) randomly adjusting the contrast of training scans by raising each scan's voxels to $\gamma$, where $\gamma = e^{\beta}$, $\beta \sim \mathcal{U}(-a, a)$, and a is set to 0.3. Volumes are presented on a normalised basis, where volumes are normalised by the mean of all volumes across the range of conditional parameters, independently for each ROI and method. In addition, a 95% confidence interval is also presented for the empirical estimates made from training data within 2-year age bins.*

#### S14 Spatial distributions of topological inconsistencies and structural similarity

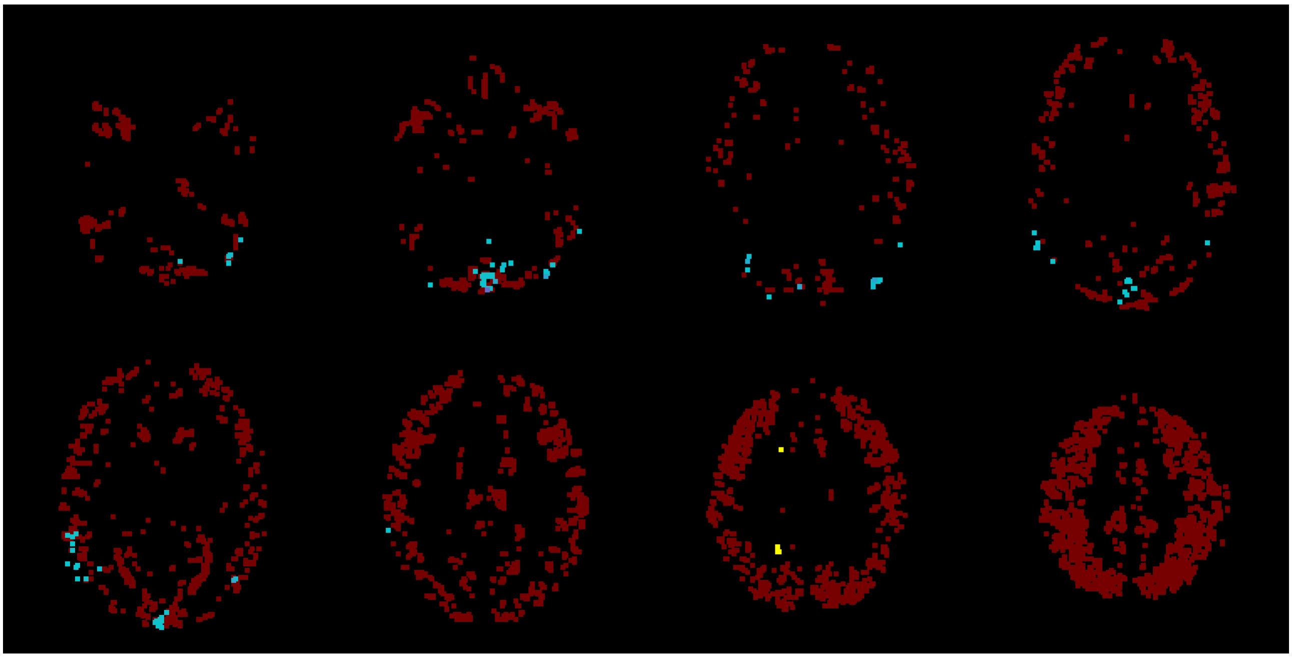

**Figure 14:** *Spatial map of negative/zero Jacobian determinants produced by the methods of Dalca et. al. (red), Yu et. al. (blue) and Dey et. al. (yellow) on the test dataset. This map is presented as a lightbox view where there are negative Jacobian determinants.*

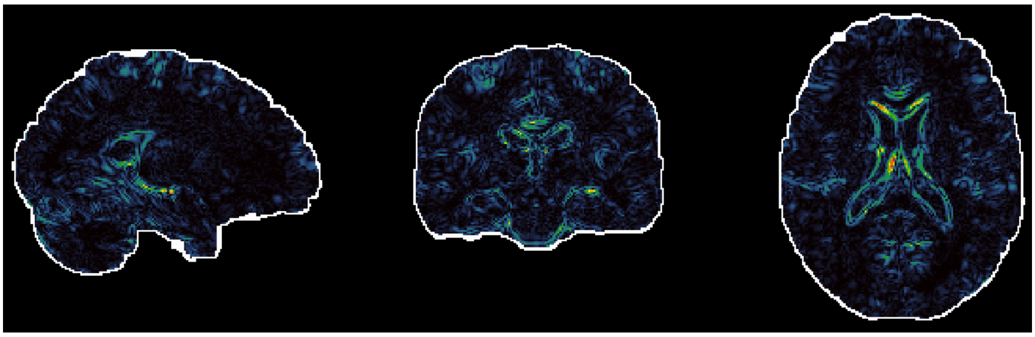

**(a)** *Ours*

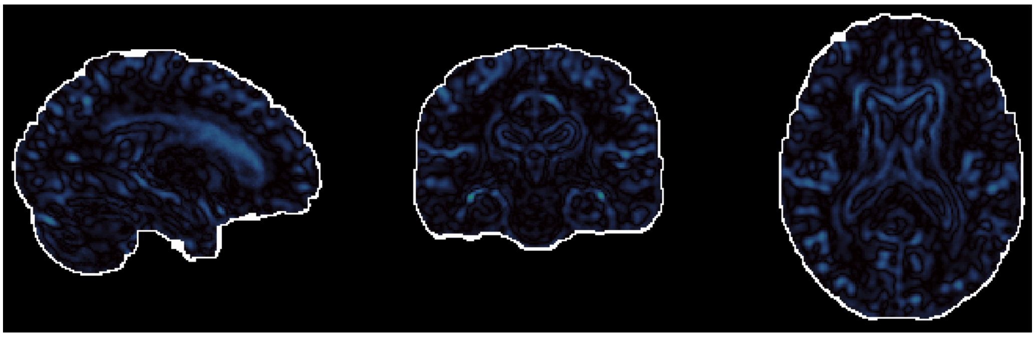

**(b)** *Dalca et. al.*

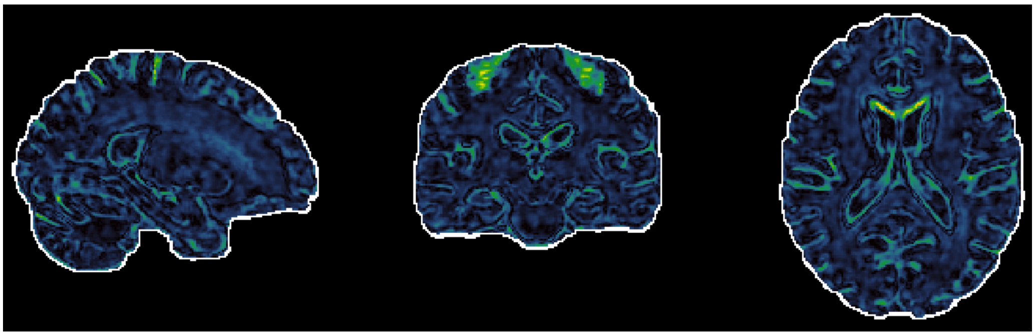

**(c)** *Dey et. al.*

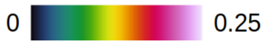


**Figure 15:** *Example spatial maps of the absolute difference between a 56-year-old conditional template and a 61-year-old conditional template (after registration of the former to the latter) for our method and the methods of Dalca et. al. and Dey et. al.. This absolute difference is shown by the colourbar presented below the maps, with range: (0, 0.25].*

#### S15 Conditional template vs unconditional template registration performance

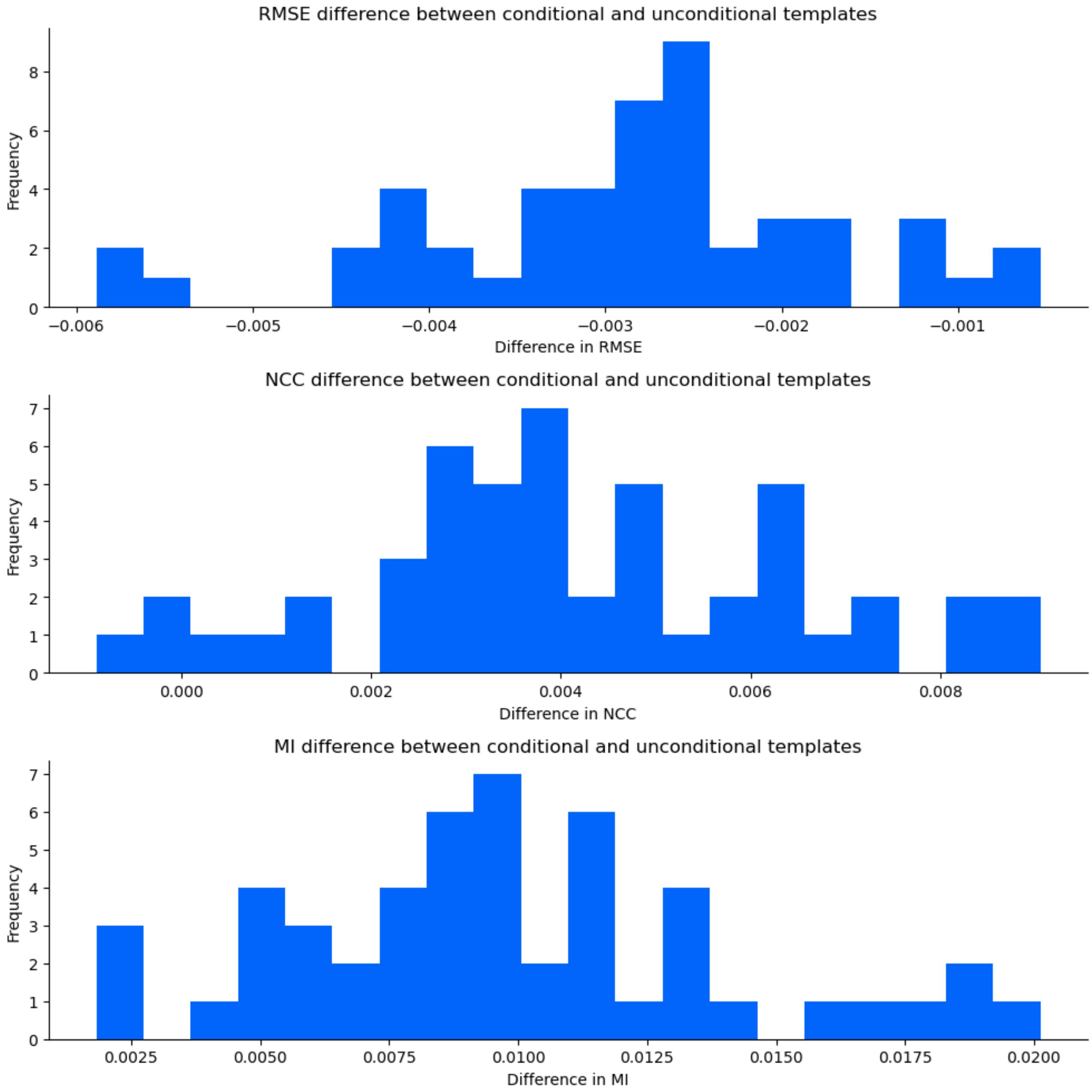


**Figure 16:** *Distributions of differences between root mean squared error (RMSE), normalised cross-correlation (NCC), and mutual information (MI) for conditional versus unconditional templates after registration to subject scans. Each difference value is calculated as the relevant metric value from a conditional template registered to a subject scan minus the metric value from the unconditional template registered to that same subject scan; i.e., we display the histograms of conditional less unconditional registration results.*

#### S16 Participant scan alignment to conditional templates

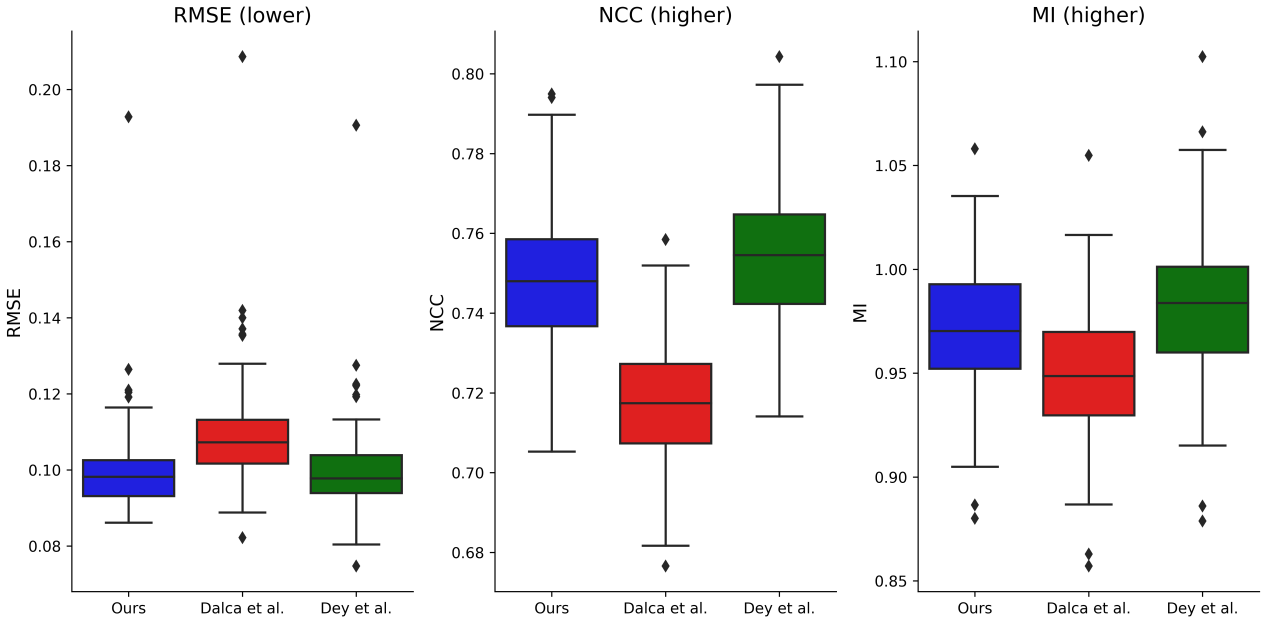


**Figure 17:** *Individual participant alignment to conditional templates across methods. We have non-linearly registered each participant scan in the test dataset to conditional templates (produced for the age of the participant at scan date using our method, as well as the methods of Dalca et al. and Dey et al.) using the SyN algorithm. RMSE, NCC and MI are calculated between each registered participant scan and its corresponding conditional template.*
